# Flexible Working Memory in the Peripheral Nervous System

**DOI:** 10.1101/2025.09.26.678884

**Authors:** Sihan Yang, Yueying Dong, Anastasia Kiyonaga

## Abstract

Working memory (WM) representations that are distributed across the brain can be flexibly recruited to best guide behavior^1–4^. For instance, information may be represented relatively more strongly in visual cortex when a WM task requires fine visual detail, or more strongly in motor cortex when a specific response can be prepared^5–10^. If WM drives goal-oriented actions, we might also expect such task-dependent signals to propagate to the peripheral effectors that realize WM commands. Accordingly, oculomotor signatures like gaze biases can track simple visuo-spatial WM features^11,12^, but their functional flexibility is unclear. Here, we test the idea that WM content is adaptively distributed across the nervous system according to behavioral demands. We ask whether patterns in both eye and hand movements can express visual WM stimulus features, and whether the distribution of such activity shifts with the task context. In a delayed recall task, we manipulated how human participants reported their memory: they would either draw a line or adjust a visible response wheel to match a remembered orientation. Via continuous eye- and stylus-tracking, we found that remembered orientations were decodable from small inflections in both gaze and hand movements during the WM maintenance period. Moreover, this decoding strength varied by response format. Gaze patterns tracked memorized features relatively better in the *wheel* condition (vs. *draw*), while hand movements were better in the *draw* condition (vs. *wheel*). Therefore, visually encoded WM contents may be adaptively allocated to task-relevant motor effectors, balancing WM representations across peripheral activity according to behavioral needs.

## RESULTS

The crux of working memory (WM) lies in its flexibility to apply mental representations toward myriad current goals. Likewise, WM content may be held in different neural response patterns depending on how it will be used^4,6,9,13,14^. Rather than sustain a precise copy of encoded sensory features, an adaptive WM code may incorporate critical features of the task context^15^. Therefore, while the quality of visual WM is often tested by having participants adjust a visible prompt to match their memory (e.g., click locations on a wheel, continuously adjust the angle of a bar)^16–19^, this test format may shape the memory itself. Here, to examine how motor response demands influence visual WM maintenance, we had participants report their memory using a digital stylus and tablet. The critical manipulation was whether they were asked to freely recall the remembered stimulus by drawing it from memory, or whether they used a more conventional response wheel to make a continuous adjustment report. We recorded both hand and eye movements throughout the task, to test whether WM content would be flexibly expressed across relevant motor effectors.

### Drawing enables precise free recall from working memory

Participants (n=35) completed a delayed report task (**Figure 1a**). They saw two sequentially presented Gabor patches and were cued to remember the orientation of either one or both patches over a delay of several seconds. They then reported the remembered angle(s) of cued stimuli by either drawing a line (in *draw* blocks; **Figure 1b-c**) or using the stylus to adjust a response wheel (in *wheel* blocks; see ***<u>Procedure and task design</u>*** for all conditions).

**Figure 1:**
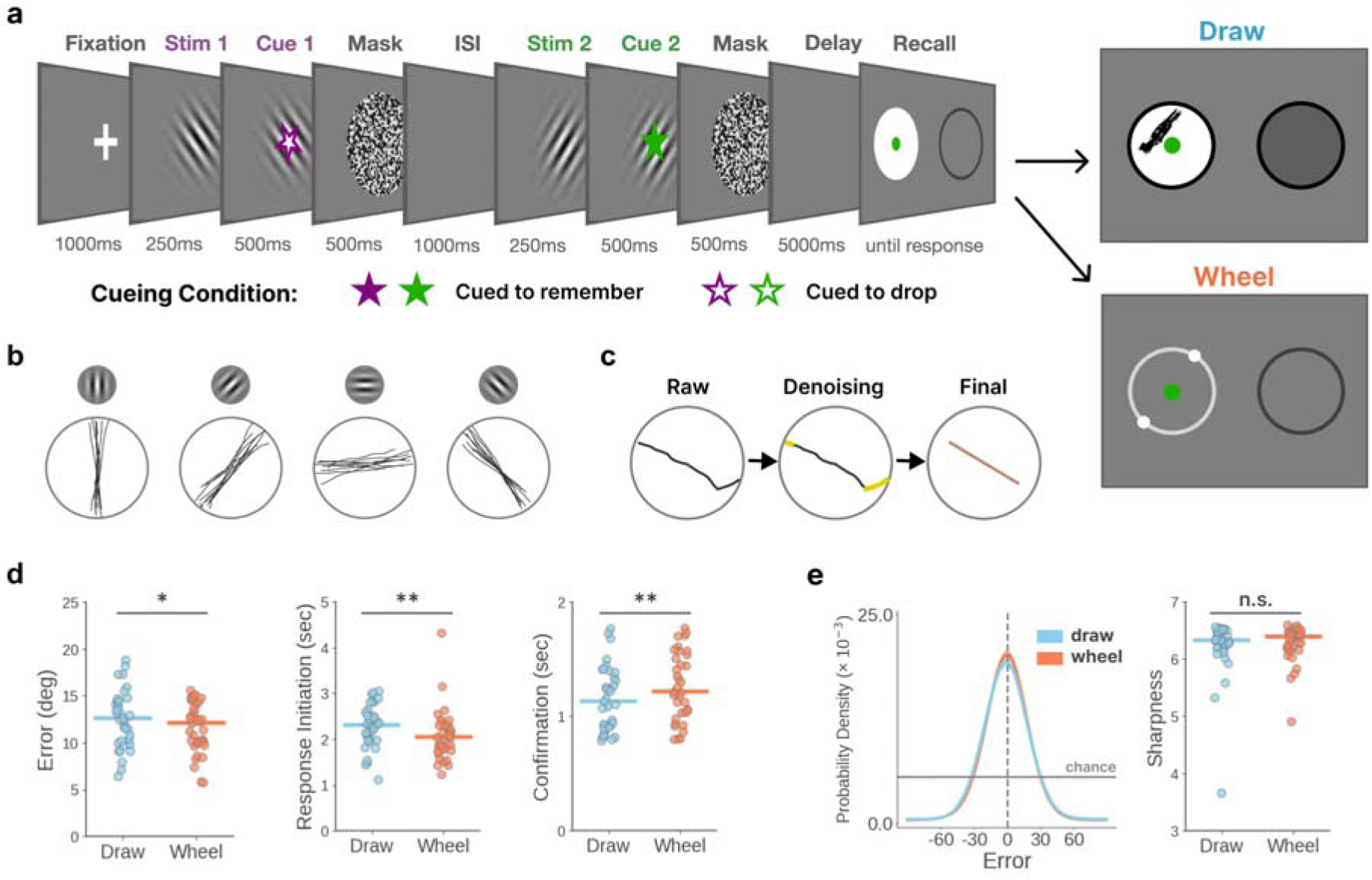
Experimental design and recall performance. **(a)** On each trial, WM sample stimuli were two sequentially-presented Gabor patches, randomly varying in orientation. A colored cue indicated whether to remember (filled star) or drop (empty star) each stimulus. Either one or both stimuli could be cued to remember, but cues appeared after stimulus onset so that both stimuli would be initially encoded. In the test phase, two circular response regions appeared, and a colored dot at the center of a region indicated which item should be recalled there. In addition to the color cue, the shading of the region indicated if it was editable: only an illuminated wheel or input field could accept a response. In both cases, participants used a stylus and tablet to respond. In the ‘draw’ condition, participants freely drew a line to report the remembered orientation inside the input field. In the ‘wheel’ condition, participants adjusted the location of two dots – appearing opposite each other along the circumference of a circle – that represented the endpoints of an oriented line. Participants could tap or drag the stylus along the wheel to adjust the dots, where adjusting one dot also moved the other. On all trials they confirmed their response by tapping a ‘continue’ button to advance. The draw or wheel response formats were instructed at the beginning of each block, which were given in quasi-random order (see **<u>Procedure and task design</u>**). **(b)** Example drawings for a range of stimulus orientations (±1°). **(c)** Each raw drawing was first denoised by removing extremities at the beginning and end of the response period (shown in yellow in the ‘denoising’ panel). Response angle was calculated as the average of the remaining strokes (‘final’ panel). **(d)** Recall performance for both draw (blue) and wheel (red) conditions, showing response error (i.e., angular difference between the response and the correct stimulus angle), response initiation time (i.e., time to first stylus contact in the response field), and response confirmation time (i.e., time from completing response to pressing the “continue” button). Point clouds show individual subject means; horizontal lines mark the group median. Asterisks mark significant Wilcoxon signed rank tests. **(e)** Smoothed distributions of response errors for the draw and wheel conditions across the group (left), and point clouds showing the sharpness of the fitted curve for each individual (right).

We tested for performance differences between *draw* and *wheel* conditions using both *t*-tests and Wilcoxon signed ranks tests. Both tests showed similar results (see ***Data S1.1***), but we report the latter in the main text, because several conditions did not meet normality assumptions for *t*-tests.

Response errors were overall larger in the *draw* condition than the *wheel* condition (***W* = 174, *n* = 35, *p* = 0. 020**; **Figure 1d**, left). Responses were also initiated more slowly in the *draw* condition (vs. *wheel*; ***W* = 125, *n* = 35, *p* = 0. 001**; **Figure 1d**, middle), but were confirmed more quickly after completion (***W* = 138, *n* = 35, *p* = 0. 003**; **Figure 1d** right). We also examined precision as the sharpness of each participant’s response error distribution (see ***<u>Recall response processing</u>***), which did not differ between *draw* and *wheel* conditions (***W* = 274, *n* = 35, *p* = 0. 512; Figure 1e**). Errors were overall larger when remembering two stimuli (vs. one), but this load effect did not interact with the *draw* vs. *wheel* response format (see ***Supplemental Results 1***), so we collapse across load conditions for most analyses.

In sum, *draw* responses showed slightly larger errors, but were overall highly precise and did not differ in sharpness from *wheel* responses. Drawing is therefore an effective tool to capture WM quality^20,21^. However, small differences in motoric response demands may influence WM performance. Namely, *draw* responses were initiated more slowly, but were finalized faster than *wheel* responses. Both response types involved precise motor control over the stylus, but drawing required articulating the whole line, while wheel adjustment required a smaller range of motion around a specific position. Subjects may have taken longer to prepare the relatively elaborated motor plan for a drawing, but then had more momentum or confidence to confirm it after completion. These behavioral differences could also stem from the different memory demands inherent to each probe. Drawing may index true free recall, because it is executed without an informative prompt^22^, while wheel responses may be subtly shaped by their dependence on a visible cue^23^. Growing evidence suggests that such task context factors might influence neural WM content representations^13,14^, and below we will examine whether they shape peripheral motor activity during the WM delay.

### Gaze and hand motion patterns maintain WM features

WM is tightly intertwined with action, and neural signatures of WM content can often reflect features of an impending motor response^8,9,24–27^. Simple features of remembered information can also now be detected in oculomotor signatures outside the brain, like gaze biases that track the remembered location or orientation of a relevant WM stimulus^11,12,28,29^. WM may therefore engage distributed structures across the nervous system that are relevant for current goals. If so, in addition to oculomotor activity, we would expect manual motor effectors to be modulated by corresponding WM demands.

Here, we tested whether memorized orientation features could be detected in both gaze and hand movement patterns during a WM delay (first collapsed across the *draw/wheel* condition; **Figure 2**). From eye movement recordings, we extracted distributions of gaze offset angle relative to each individual’s baseline gaze location (**Figure 2a**); From stylus recordings, we extracted the distribution of movements along each direction (**Figure 2b**; see ***<u>Quantification and Statistical Analysis</u>***). During the blank WM delay interval, we observed that small inflections in both gaze offsets and hand trajectories appeared to vary systematically with the memorized orientations (**Figure 2a-b**), suggesting that motor activity tracks WM content information. We next formally quantified these patterns.

**Figure 2:**
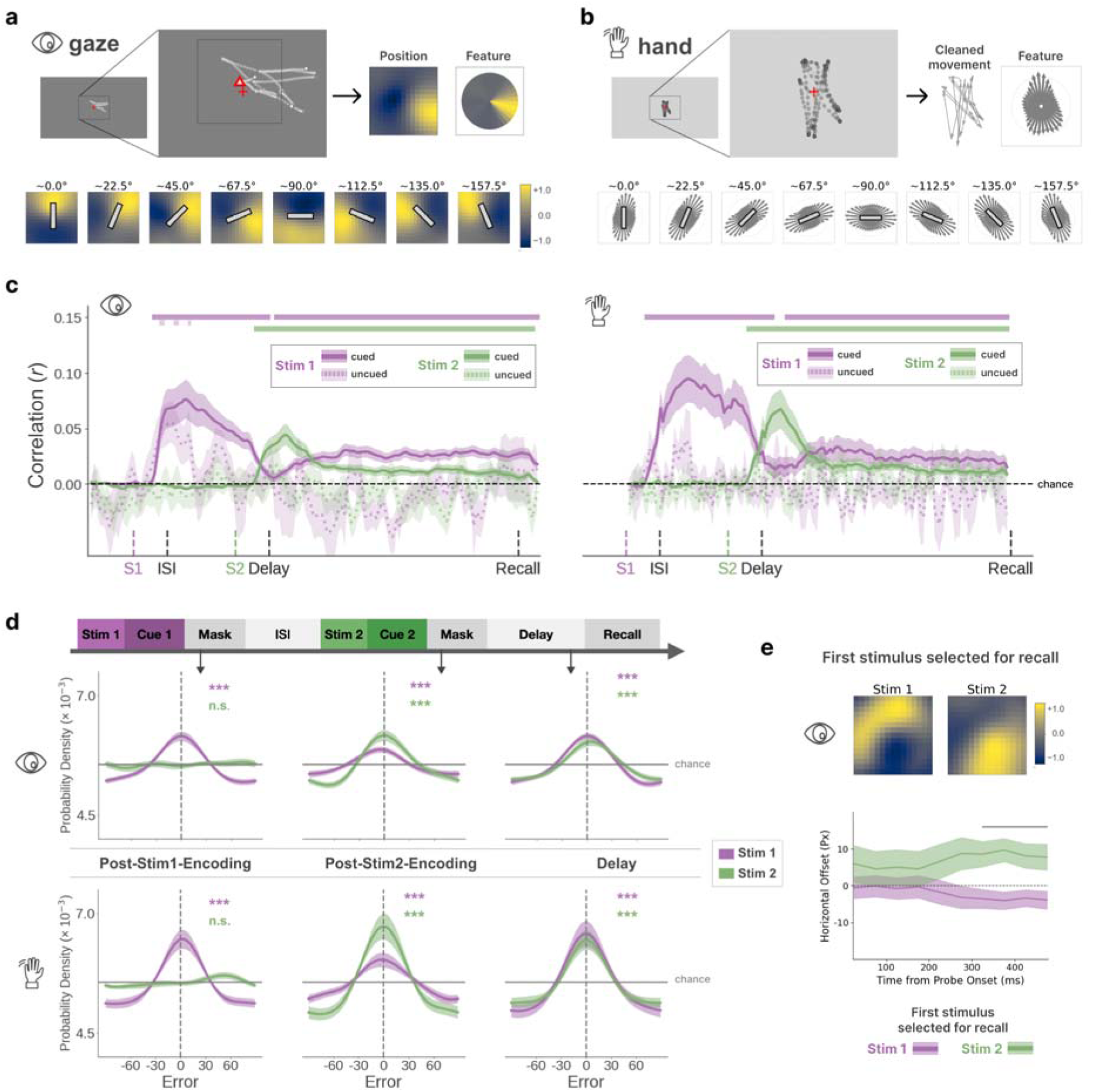
Orientation information in gaze and hand motion patterns during working memory. **(a)** Top: Each participant’s median gaze location served as baseline (denoted by red triangle). After baseline subtraction, we computed distributions of gaze offset angle. Bottom: Normalized 2D heatmaps of gaze positions during the delay, aggregated across participants and conditions. Each central bar reflects the bin center for a given subset of trials, split into 8 orientation bins. Yellower color indicates more visitations at that location. **(b)** Top: Example stylus movements for computing each participant’s angular distribution. Bottom: median-centered angular distributions of hand movements during the delay, aggregated across participants and conditions. The length of each vector is proportional to the magnitude of movements in that direction. Each central bar reflects the bin center orientation for that subset of trials. **(c)** Timecourse of representational similarity for gaze (left) and hand motion (right) patterns, collapsed across load 1 and 2 trials. The y-axis shows the strength of correlation between the motor activity and stimulus similarity matrices. Bold (dashed) traces show correlations for stimuli that were cued to remember (drop). Error bands reflect standard error of the mean. Shaded horizontal bars mark time periods when the correlation was significantly different from chance (cluster-based permutation test, p < .05). **(d)** Inverted encoding modeling (IEM) of stimulus orientation, for cued items, during key trial epochs. After training, the model predicts the remembered orientation on each trial as a probability distribution from 0° to 180°. Distributions were recentered around 0° orientation (clockwise positive), and each plot depicts the distribution of model errors (i.e., the ‘reconstruction’), averaged across trials and participants. Reconstructions from a leave-one-subject out model are shown for both gaze (top) and hand motion (bottom) data. Asterisks denote whether the sharpness is significantly greater than expected from a uniform distribution (i.e., zero). **(e)** Gaze biases during WM recall. Trials were grouped according to whether subjects first reported Stim1 or Stim2. Top: Gaze distributions aggregated across the 500ms after probe onset. Bottom: mean horizontal gaze positions over time, when selecting Stim1 (purple) or Stim2 (green). Gray horizontal bar indicates period where the two differ significantly (cluster-based permutation test). (Icon ‘Eye’ by Natalia, ‘Hand’ by Paffi from the Noun Project (https://thenounproject.com) used under CC BY 3.0.)

To gauge the quality of WM stimulus information in eye and hand movement patterns, we applied two multivariate techniques that are commonly used for neuroimaging analysis. First, we used representational similarity analysis (RSA^30^; see ***<u>Representational similarity analysis</u>***) to assess whether more similar orientations evoked more similar eye or hand movement patterns during WM^12^. This would indicate that peripheral motor signatures discriminate between remembered stimuli. To test this, we correlated the stimulus similarity matrix (pairwise similarity between stimulus orientations) with the similarity matrix for the peripheral motor signatures (pairwise cosine similarity between movement feature vectors across remembered orientations). We assessed the timecourse of this correlation across the trial (**Figure 2c**).

Shortly after encoding, stimulus-specific information emerged in both gaze and hand movement patterns for stimuli that were cued to remember (‘cued’), and it remained above chance throughout the WM delay. Critically, no above-chance information emerged for stimuli that were perceived but then cued to drop (‘uncued’). Therefore, both manual and oculomotor signatures reflect only the task-relevant mnemonic content – they are explained by neither lingering sensory-evoked activation nor a reflexive motor response to items that are seen but not remembered. This remained the case whether one or both stimuli were cued to remember, indicating that these peripheral motor patterns can reflect unique stimulus evidence for multiple items in mind (**Figure S2c**).

While RSA can tell us that activity patterns reflect the relational structure in the stimulus space, the analysis does not derive the specific item being remembered. So we also applied inverted encoding modeling (IEM^31,32^) to reconstruct an estimate of the remembered orientation from aggregate motor patterns. IEM learns stimulus-activity associations across feature-selective information channels during training, then predicts what information is reflected in untrained channel responses. The approach is used widely to estimate WM representation quality from population-level neural activity (e.g., fMRI/EEG)^33–39^, and here we apply it to peripheral motor activity to approximate the information in those patterns (see ***<u>Inverted encoding modeling</u>***). We trained and tested models using leave-one-subject-out cross-validation (*across-subject*) which we report in the main text. We report 5-fold cross-validation (*within-subject*) results in ***Supplemental Results 2.3*** (**Figure S2d**).

We were able to reliably reconstruct relevant stimulus orientations during key trial epochs (**Figure 2d**; where stimulus evidence is the sharpness of the model prediction error distribution). Right after the first sample stimulus, that orientation was reliably reconstructed (***p*< 0.001** for Stim1, gaze and hand). Right after the second stimulus, that stimulus was reliably reconstructed, whereas the first stimulus remained above chance but became relatively dampened (***p*< 0.001** for Stim1 and Stim2, gaze and hand). During the blank WM delay, both relevant stimulus orientations could be reliably reconstructed, from both gaze and hand movement patterns (all ***p* < 0.001**; see ***Data S1.2***). IEM evidence for each cued stimulus was also comparable whether one or both stimuli were remembered (i.e., load of 1 or 2; **Figure S2c**), further suggesting that these peripheral activity patterns can code for multiple memory representations.

To summarize, we used two multivariate approaches to quantify the WM information content in gaze and hand movement patterns, across encoding and maintenance intervals. We found robust evidence for task-relevant mnemonic content in both motor measurements, suggesting that visual WM content is expressed in multiple effectors during maintenance. Stimulus information within individuals could also be reconstructed from models trained on other subjects’ data, indicating that these motor WM signatures may share properties across people.

### Gaze biases during recall reflect selection from WM

If the observed motor signatures of WM are geared to support goal-directed behavior, they may also carry into the recall phase as a response is being prepared. For instance, directional gaze biases have been implicated as a marker of attentional selection within WM (i.e., in response to a retro-cue^11,40^), and they may similarly reflect selection of WM items for explicit recall.

Here, we focus on gaze during the 500ms after probe onset, for trials when both stimuli were cued, to capture the stimulus retrieval process prior to motor execution (see ***<u>Gaze biases during selection for recall</u>***). We sorted trials according to whether Stim1 or Stim2 was reported first, and compared gaze between those two conditions. We found distinct gaze biases when preparing to report Stim1 vs. Stim2, even though both stimuli were encoded at screen center and response field locations were randomized from trial-to-trial (see ***<u>Procedure and task design</u>***). Gaze was shifted overall leftward when selecting Stim1 and rightward when selecting Stim 2, and this bias grew across the selection period (**Figure 2e**). Gaze therefore appears to reflect systematic biases associated with the temporal order of stimulus encoding, which may anchor retrieval for each item^41–43^. Gaze also preferentially coded for the specific orientation of the stimulus that was being selected (vs. the other item; **Figure S2e**), highlighting that eye movement patterns can express multiple levels of WM information – both stimulus feature and temporal context – during selection for a manual response.

### Motor signatures of WM shift with expected response format

Cortical signatures of WM content can adapt to how WM will be tested^3–6,14^. For instance, visuo-spatial WM content may be offloaded to a more motor-like neural representation when a specific response can be prepared^9^. This suggests that motor intentions might mold how or where visual WM content is maintained. In other words, the same visual WM content might be represented differently when it precedes different actions. Our theoretical framework predicts that WM may also flexibly recruit the peripheral nervous system according to the task context. To test this, we next asked whether the WM signals in gaze and hand movements vary when a *draw* or *wheel* recall response is required.

Both *draw* and *wheel* tasks tested memory for the same stimulus feature (angle), and used the same response implement (stylus). Nonetheless, we found that the relative strength of WM stimulus information in gaze and hand signals was modulated by the response format condition (**Figure 3**). Namely, the correlation in the gaze RSA was stronger in the *wheel* condition compared to *draw*. Conversely, the correlation in the hand motion RSA was stronger in the *draw* condition compared to *wheel* (**Figure 3a**). Therefore, we might infer that the motor effectors are differently task-relevant in the two conditions – while the eyes appear to ‘prefer’ the *wheel* condition, the hands appear to prefer the *draw* condition.

**Figure 3:**
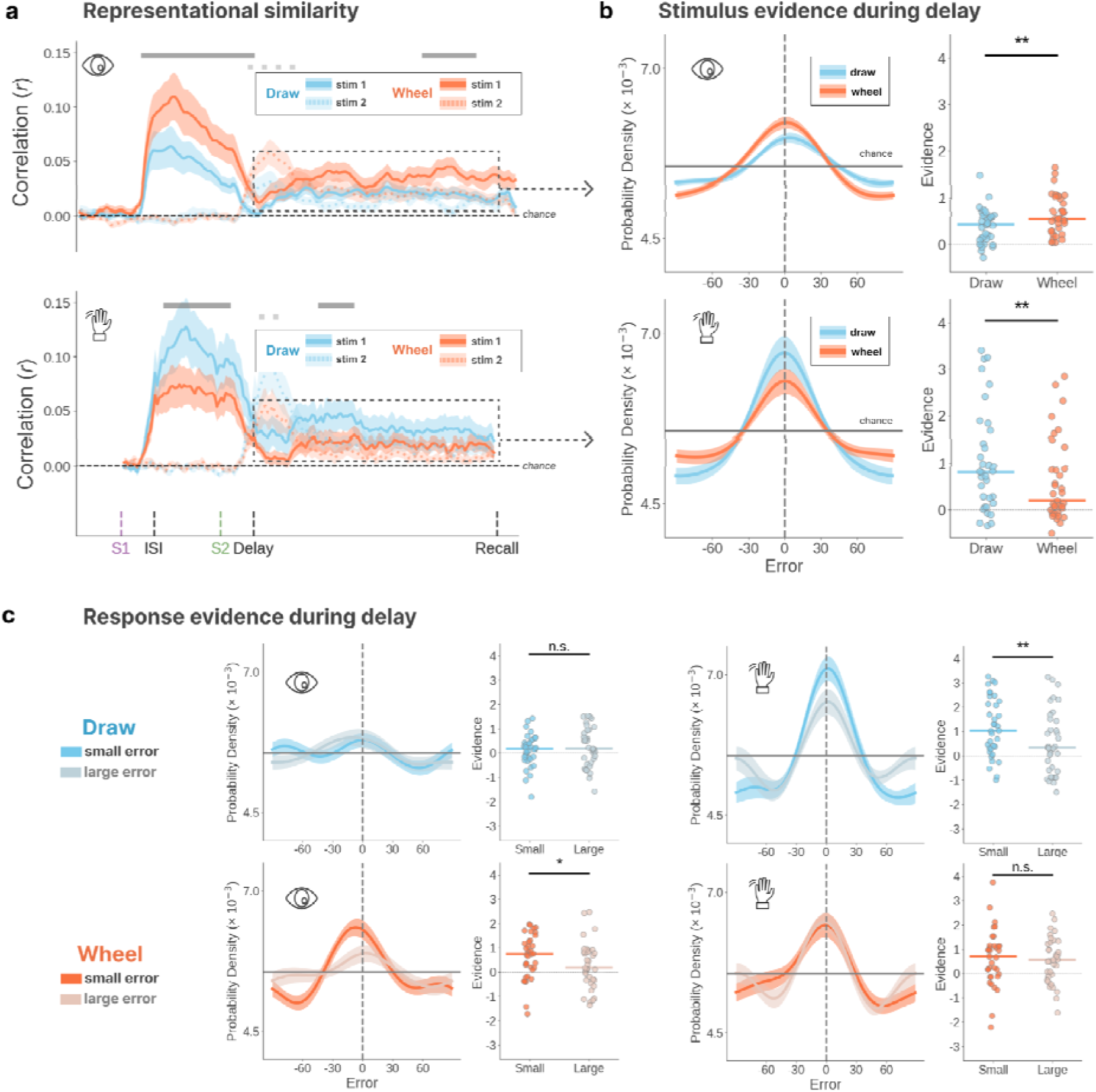
WM representations in gaze and hand motion under different response formats. **(a)** Time course of representational similarity for gaze data (top) and hand motion data (bottom), comparing the draw (blue) and wheel (red) conditions. The solid (dashed) grey bars mark time points where the correlation for Stim1 (Stim2) differed between conditions (cluster-based permutation test, p < .05). **(b)** Left: IEM reconstructions of remembered orientations, during the delay period, for draw and wheel conditions (combined across Stim1 and Stim2). Right: Points represent each individual’s IEM-based WM stimulus evidence (i.e., the sharpness of the error distributions), and colored horizontal lines show the group median for each condition. Asterisks denote whether the evidence differed significantly between draw and wheel conditions. **(c)** Delay-period IEM reconstructions and individual-level evidence for analyses conditioned on the upcoming response (rather than the encoded sample stimulus). After testing, each subject’s responses were sorted based on error magnitude, and here response evidence is displayed for each subject’s top and bottom error tercile. E.g., ‘small error’ conveys highest precision responses, while ‘large error’ conveys lowest precision responses (restricted to trials of load=2). Asterisks denote whether the evidence was significantly greater for ‘small error’ responses than ‘large error’ ones.

This apparent preference was also evident in the IEM-based reconstructions from the delay period (**Figure 3b**). Gaze-based reconstructions showed stronger stimulus evidence (for both Stim1 and Stim2) in the *wheel* condition (vs. *draw;* ***W* = 154, *n* = 35, *p* = 0. 007**). Conversely, hand-based reconstructions showed stronger stimulus evidence in the *draw* condition (vs. *wheel*; ***W* = 132, *n* = 35, *p* = 0. 002**; **Data S1.3**). This difference between *draw* and *wheel* response formats also held on trials when both stimuli were remembered (**Figure S3d**), indicating that the effect is reliable across WM loads.

These analyses show that, during a WM delay, upcoming response demands may modify how information is represented. Relatively stronger hand-based representations during the *draw* condition (vs. *wheel*) may reflect the need to convert stimulus information into a precise manual action plan. Relatively stronger gaze-based representations during the *wheel* condition (vs. *draw*) could reflect the greater need to rely on a visual representation for stimulus comparison. The WM feature information expressed in peripheral motor signatures may strategically adapt to expected task demands.

### Effector-specific WM activity precedes precise responding

The effector-specific preferences for *draw* vs. *wheel* conditions also emerged in analyses that were conditioned on the upcoming response, rather than the encoded sample stimulus (i.e., models trained and tested on the angle that was ultimately reported; see ***<u>Inverted encoding modeling</u>***; **Figure 3c**). These analyses can illuminate whether delay activity relates to motor preparation or WM quality. If purely motor preparation, the patterns should predict the upcoming response equally well for both low- and high-error recalls. Instead, response evidence was stronger preceding low error responses (vs. high error), specifically in the ‘preferred’ effector for the task condition (gaze/wheel: ***W* = 429, *n* = 35, *p* = 0. 031**; hand/draw: ***W* = 488, *n* = 35, *p* = 0. 002**; **Data S1.4**). This illustrates that preferential eye (hand) engagement during the *wheel* (*draw)* condition precedes more precise recall.

Note, however, that the differences between *draw* and *wheel* conditions likely reflect quantitative differences in representational strength rather than qualitative differences in coding format. While hand movements were somewhat more frequent and pronounced in the *draw* condition overall (**Figure S3a**), data from the *draw* condition could be used to successfully reconstruct WM content in the *wheel* condition (and vice versa; **Figure S3c**), suggesting shared coding properties between response formats (**Figure S3b**).

### Competition and cooperation between gaze and hand codes

If the different response demands modulate which effectors are most relevant for WM maintenance, we might expect to observe a hand-eye trade-off in stimulus evidence between the *draw* and *wheel* conditions (**Figure 4a**). For instance, when the response format changes from *wheel* to *draw*, a greater increase in an individual’s hand evidence should be accompanied by a corresponding decrease in gaze evidence. To test this, we calculated the difference in IEM evidence between the *draw* and *wheel* conditions, separately for gaze and hand data, for each individual (**Figure 4b**). We then examined the relationship between the gaze and hand difference scores (linear regression).

**Figure 4:**
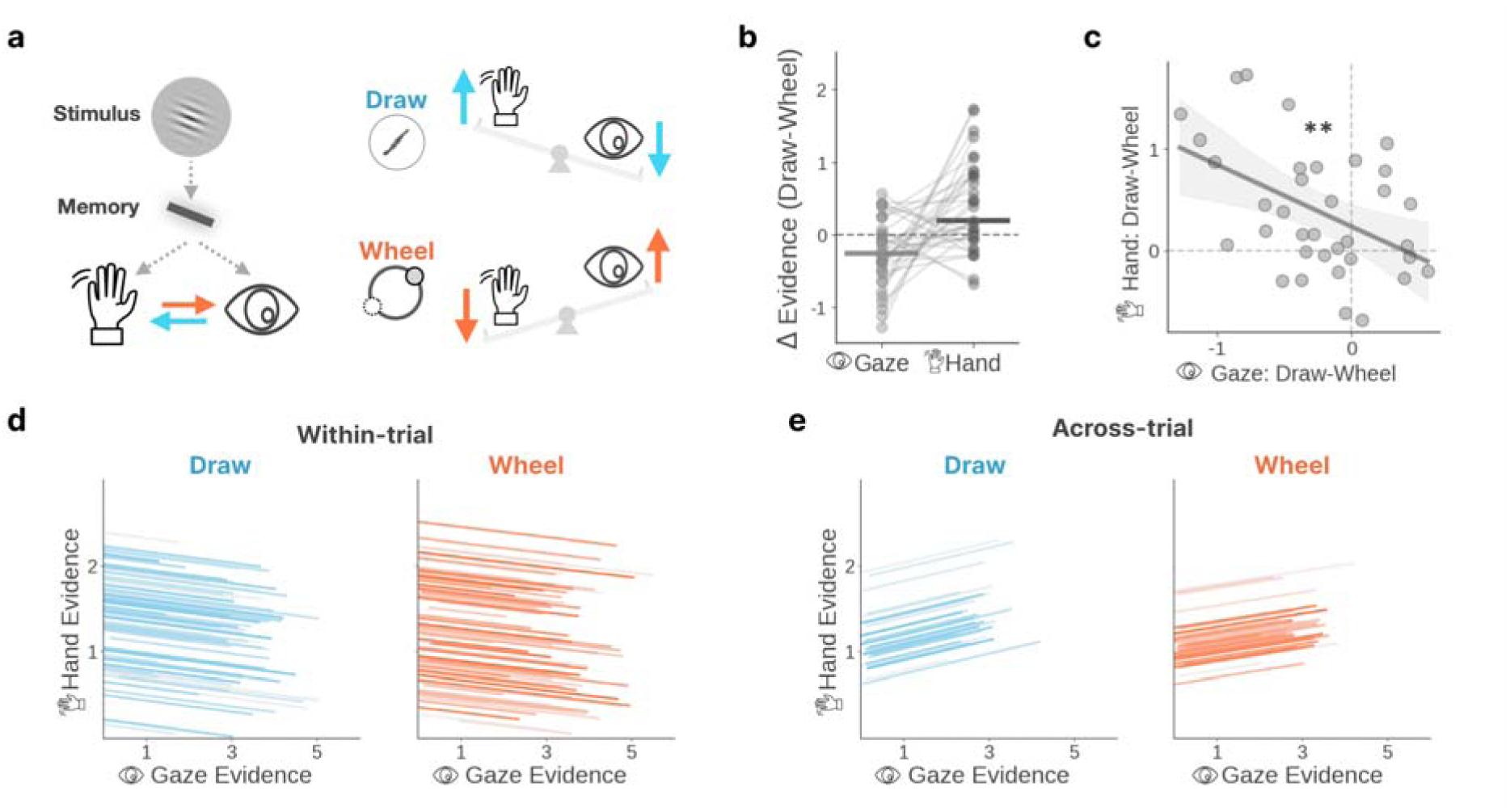
Coupling between gaze and hand-movement WM codes. **(a)** Schematic depiction of the current framework: WM is adaptively balanced across motor effectors based on task demands. The balance favors hand activity when having to draw, and gaze activity when using a wheel response. **(b)** The difference in IEM evidence between the draw and wheel conditions (draw - wheel) for each subject, computed separately for gaze and hand data (aggregated across the delay). The lighter (gaze) and darker (hand) points represent the differences between the blue and red points in the upper and lower panels of Figure 3b, respectively. A positive (negative) difference score indicates stronger stimulus evidence in the draw (wheel) condition. **(c)** Scatterplot shows the relationship between difference scores in gaze and hand movement patterns, where each point represents an individual subject. **(d-e)** Within-subjects variation in representational strength, based on IEM evidence from a 350ms sliding window across the delay. The maximum stimulus evidence (for either Stim1 or Stim2) was taken as the representational strength for that window. **(d)** Repeated measures correlations showed negative relationships between the timeseries of gaze and hand representational strength within trials. Since this correlation was conducted within each individual, the effect is shown for one example subject, where each line represents a trial. **(e)** Repeated measures correlations showed positive relationships between gaze and hand representational strength across trials. For each trial, representational strength for each modality was averaged over the timeseries, and each line represents an individual subject’s correlation across trials.

We found a negative relationship across subjects (***β* = -0. 47, *p* = 0. 005**; **Figure 4c**), consistent with the idea that the gaze and hand WM representations trade off with each other. For instance, individuals with a relatively better average hand-based reconstruction during *draw* (vs. *wheel*) also showed a relatively worse gaze-based reconstruction during *draw* (vs. *wheel*). This suggests that when the balance of maintenance activity shifts towards the hands, it is correspondingly dampened in the eyes (and vice versa). The system may adaptively distribute limited maintenance resources across the relevant peripheral structures.

If maintenance activity is apportioned across effectors, they may further trade-off with each other within a condition and individual. Lastly, we examined the correlated fluctuations between eye and hand representations within subjects, both within and across trials of the same type. We calculated a time-series of ‘representational strength’ across the delay, for each individual and trial (see ***<u>Correlated evidence fluctuations across effectors</u>***; **Figure S4a**), and we tested the correlations between the gaze and hand time-series. This relationship was negative within *wheel* condition trials (***r*** = -**0. 01**, ***W*** = **167**, ***n*** = **35**, ***p*** = **0. 014**), and descriptively negative within *draw* condition trials (***r*** = -**0. 01**, ***W*** = **200**, ***n*** = **35**, ***p*** = **0. 060**; **Figure 4d**). In other words, at time points when ‘representational strength’ was relatively greater in gaze evidence, it was relatively weaker in hand evidence — further supporting the idea that gaze and hand activity may trade-off within an individual. However, the trial-wise averages of this representational strength were also positively correlated between gaze and hand activity across trials (*draw*: ***r* = 0. 15, *p* < 0.001**; *wheel*: ***r* = 0. 11, *p* < 0.001**; **Figure 4e**), suggesting that trials with stronger baseline information in one effector also exhibited stronger information in the other.

WM codes in gaze and hand activity may be complementary. The effectors can trade-off with each other between trial conditions, and at each time-point within a trial. But gaze and hand activity may co-vary together across trials of the same type, consistent with their tight linkage in eye-hand coordination^44–47^. WM activity in eyes and hands may share a central source, yielding joint engagement during visuo-motor WM.

## DISCUSSION

Here, we found that patterns in both eye and hand movements can express visual WM stimulus features, but the distribution of such peripheral motor activity shifts with the task context. Given identical visual stimulus content, distributed WM maintenance activity may tailor to which effectors are most relevant for an upcoming task.

The hands showed stronger WM information preceding a *drawing* response (vs. *wheel*), while the eyes showed stronger WM information preceding a *wheel* adjustment (vs. *draw*). A *draw* response can be freely executed without visual input, but may require a relatively elaborated motor plan. A *wheel* response is prompted by a visible probe, and may require continuously adjusting the probe to match memory. These subtle differences in impending task demand may shape how WM activity is distributed across the nervous system.

These findings echo work showing that cortical WM signals can flexibly reflect various motor and task factors^6,8,9,27,48^. Here, we show that WM may similarly recruit the peripheral nervous system in a widely distributed and task-flexible scheme. Likewise, mounting evidence indicates that cognitive computations can be detected in bodily movements and peripheral signals, which may also adapt to expected demands^28,49–57^. These signatures have been construed as an accessible read-out of underlying neural signals^58^, but they could potentially serve a more adaptive functional role. For instance, even small movements might constitute a form of covert cognitive offloading^59^, where some of the maintenance burden is allocated to specialized circuits. This might compress the representation to the action-critical dimensions and minimize interference in central processing. Alternatively, the peripheral movements might reinforce the cortical representations in a reciprocal feedback loop. For instance, cognitive eye movement and pupil size modulations can shape visual cortex representations^60–62^, and hand gestures can improve children’s learning^63^. An intriguing possibility is that peripheral WM signatures may be more than a reflection but an active component of the distributed WM representation.

WM is increasingly understood to be intertwined with goal-oriented actions^24,25,64–68^, thus WM maintenance representations can be confounded with those for response preparation^26^. However, the signals detected here may be considered unique from simple response preparation for several reasons. For one, eye movements do not directly map onto a task-relevant response here. And although hand movements could in theory rehearse an upcoming action, the specific item to recall was often unpredictable. Moreover, delay period activity patterns were more predictive of precise recall than of the submitted response, per se. Nonetheless, if a core function of WM is to prepare for action, then even mere response preparation may be considered a meaningful aspect of the representation.

Together, these findings prompt inquiry into when and why individuals vary in engaging the peripheral nervous system during WM, and whether these signatures play a functional role. Future work could address whether this activity shoulders some of the burden for cortical storage mechanisms, reciprocally fortifies cortical signals, or reflects behavioral implementation of central representations.

## RESOURCE AVAILABILITY

### Lead contact

Requests for further information and resources should be directed to and will be fulfilled by the lead contact, Sihan Yang.

### Materials availability

No new materials were generated in this study.

### Data and code availability

● Data have been deposited to the Open Science Framework (OSF) and will be publicly available upon publication.
● All original code has been deposited to the Open Science Framework (OSF) and will be publicly available with a permanent DOI upon publication.
● Any additional information required to reanalyze the data reported in this paper is available from the lead contact upon request.

## ACKNOWLEDGMENTS

We thank Connie Xie, Yun-Chen Hung, Mycah Gutierrez and Lana Gaspariani for their help with data collection. We also thank Pria Daniel, Ana Chkhaidze and Zhuojun Ying for helpful discussions and feedback throughout this study. This work was funded in part by NIH award 1R01EY036843-01 to A.K.

## AUTHOR CONTRIBUTIONS

Conceptualization, S.Y. and A.K.; Data Curation, S.Y. and Y.D.; Formal Analysis: S.Y.; Funding Acquisition, A.K.; Investigation: S.Y. and Y.D.; Methodology, S.Y., Y.D, and A.K.; Resources, S.Y. and Y.D.; Software, S.Y. and Y.D.; Supervision, A.K.; Validation, S.Y. and A.K.; Visualization, S.Y.; Writing – original draft, S.Y. and A.K.; Writing – review & editing, S.Y., Y.D, and A.K.

## SUPPLEMENTAL INFORMATION

Document S1. Figures S1–S4, Tables S1-S4, and supplemental references.

Data S1.

## STAR METHODS

### EXPERIMENTAL MODEL AND STUDY PARTICIPANT DETAILS

A total of 45 human participants were recruited through the SONA participant pool at the University of California, San Diego (UCSD). Participants received course credit as compensation. The study was approved by the UCSD Office of IRB Administration (OIA), and all participants provided informed consent electronically before participating.

Among them, 37 successfully completed the experiment without interruption. One participant was excluded from analyses due to failing eye-data quality control on more than 40% trials, and one other participant excluded for no hand movements detected in more than 40% trials. This resulted in a final sample of 35 participants (28 female, 6 male, 1 non-binary; age: 18 to 26, M=20.50, SD=1.77, 2 declined to report). Sex, race, and ethnicity were self-reported at enrollment. Participants identified as Asian (n=14), White (n=10), multi-racial (n=6), and declined to report (n=5). 10 participants identified as Hispanic or Latino. Gender identity, ancestry, and socioeconomic status were not collected. Sex-based analyses were not conducted, which may limit the generalizability of findings.

One participant was left-handed. All participants had normal or corrected-to-normal vision (with 11 using contact lenses), and reported no history of neurological or psychiatric disorders.

### METHOD DETAILS

#### Procedure and task design

Participants sat in a dimly lit room, approximately 60 cm away from a 24-inch monitor with a resolution of 1920 × 1080 pixels. A chin rest was used to minimize head movements and maintain a consistent viewing distance. A tablet was placed in front of the monitor, on the side of the participant’s dominant hand. Participants were allowed to adjust the tablet’s position for comfort during the calibration phase, but were instructed not to move it during the experiment.

The experiment was programmed and presented using Psychopy (v2024.2.4^69^) on a Windows 10 system. Each trial began with a 1000ms fixation period, followed by two sequentially-presented Gabor patches (referred to as ‘Stim1’ and ‘Stim2’ respectively) on a neutral grey background. The orientation of each patch was independently sampled from a uniform distribution ranging from 0° to 180°. Each Gabor had a Gaussian envelope with a radius of 0.16 screen-heights (≈4.5° visual angle) and a spatial frequency of 5 cycles per screen height (≈0.18 cycles/deg, yielding ∼1.6 cycles across the patch).

A cue appeared 250 ms after each Gabor patch onset, in the form of a star overlaid atop the center of the Gabor, for 500ms. The color of the star tagged the stimulus so that it could be cued later for recall, and the filling of the star indicated whether that stimulus should be remembered (filled star) or could be dropped (hollow star). The color-cue mapping (i.e., which color corresponded to Stim1 and Stim2) was randomized across trials.

Each Gabor stimulus and its cue offset simultaneously, followed by a noise mask. The mask consisted of random black (RGB: 0,0,0) and dark-gray (RGB: 187,187,187) squares, with a mean luminance of ∼93 (on 0–255 scale) and a spatial frequency of ≈5.6 cycles/deg, filling the same area as the Gabor stimuli. The first mask (after Stim1) was followed by a 1s blank delay, referred to as the inter-stimulus interval (ISI). The second mask (after stim2) was followed by a longer blank delay of 5s. This latter delay is the WM delay period on which IEM analyses are focused.

During the recall phase, participants reported their memory of the cued orientation(s) using a stylus and tablet. The response fields and cursor became visible immediately at recall phase onset, and the stylus worked like a mouse. Moving the stylus above the tablet moved a visible cursor, and touching down on the tablet was akin to clicking. In the *draw* condition, participants were instructed to draw a single line to best match the angle of the remembered orientation. A white circular input region appeared, and participants could draw anywhere within the region. In the *wheel* condition, participants adjusted the positions of two opposing dots on a response wheel. A bare light grey response wheel (a plain circle) appeared initially. When the stylus first touched down on the tablet, within range of the wheel, one dot appeared at that wheel location and a corresponding dot at the opposite location. The two dots formed an imaginary line, and the subject’s task was to adjust the dots so that the line matched the remembered orientation.

The response phase was self-paced, and participants were able to modify their responses until they were satisfied to press a ‘continue’ button at screen center. To adjust a *draw* response, participants could use a ‘redo’ button below the response field to erase the drawing and make a new one. To adjust a *wheel* response, participants could either tap a new wheel location with the stylus or drag one dot along the wheel circumference to adjust its position, and adjusting the position of one dot automatically adjusted the other.

To prevent anticipatory gaze or hand motion biases to expected locations, the positions of the response fields were randomized across trials. The two regions could be arranged either horizontally (to the left and right of the screen center) or vertically (above or below the screen center), with equal probability (**Figure S1f**). The two sample stimuli were also assigned randomly to the two response regions (left vs. right, up vs. down), with equal probability. During the recall phase, two response fields were always visible, but only illuminated fields were editable to receive a response. Such editable regions were also marked by a central colored dot (green or purple), indicating which stimulus should be reported within. On trials where one WM item was probed, only one response region would be illuminated, whereas both would be illuminated on trials where both items were probed (and participants could report the items in any order they chose).

The task also included some conditions that are not analyzed here. Namely, the experiment included two types of blocks: *explicit* and *neutral*. In *explicit* blocks, when both items were cued to remember, they were both probed for recall (**Figure S1a**). In *neutral* blocks, while both items were always cued, there was a 50% chance that only one of them would be probed (**Figure S1b**). Therefore, across all conditions, participants may have been cued to remember either one or two items, and sometimes they only had to recall one item even if they were cued to remember two. However, the cues were always “valid” in that participants were never tested on an item that was uncued (see Table of trial/block conditions).

| Block | Trials |
| --- | --- |
| **Explicit:** During encoding, participants were certain that any cued item(s) would be probed later. | □ 50%: Remember one, recall one |
|  | □ 50%: Remember both, recall both |
| **Neutral:** Participants were cued to remember both items, but were uncertain whether one or both of the cued items would be probed later. | □ 50%: Remember both, recall one |
|  | □ 50%: Remember both, recall both |

The experiment consisted of 16 blocks: 8 each in the *draw* and *wheel* conditions. Each block contained 10 trials, yielding 160 trials in total (80 per response format condition). The experiment was conducted in two batches separated by a 10-minute break. Each batch consisted of 8 blocks, which were either all *explicit* or all *neutral* probe conditions, with the order randomized across participants. Within each batch, the first or second set of 4 blocks was randomly assigned to the *draw* or *wheel* condition. Thus, participants completed *draw* and *wheel* conditions in alternating sets of 4, which was aimed to minimize context-switching demands. Before each batch of the experiment, participants reviewed a diagram outlining the design of the upcoming blocks (**Figure S1a** or **b**), and completed 12 practice trials to familiarize themselves with the task.

The number of trials per subject (i.e., 160 trials) was largely based on practical considerations. During piloting, this was the maximum number of trials participants could complete without fatigue and severe detriment to performance. However, as described below, the across-subjects analysis approach is designed to be robust to small trial numbers per person, per condition.

##### Gaze data recording

Gaze data were recorded using a desk-mounted EyeLink (Portable Duo) eye tracker (SR Research Ltd), with a sampling rate of 1000 Hz. The eye tracker was calibrated before each batch of the experiment, with drift correction performed before each block. On each trial, gaze data were recorded from onset of the fixation period until 500 ms after the start of the response phase. Data were exported using the EyeLink Data Viewer software package (v.4.3.1)^70^.

##### Hand motion data and recall response recording

Incidental hand motion data and explicit recall responses were both recorded using a Wacom tablet (Intuos CTH-690), which captures stylus movements with a resolution of 2540 lines per inch (LPI) and has an active area of size 8.5 x 5.3 in. The stylus functioned as a direct input device, with the tip position over the tablet mapped to the cursor position on the screen, effectively acting as a mouse. The tablet was smaller than the monitor, and a movement of 1cm on the tablet would correspond to a measurement of 90.8 pixels. Likewise, the spatial resolution of the stylus position matched the monitor resolution of 1920×1080 pixels. Therefore, we were able to capture submillimeter motor fluctuations on the order of ∼0.1-0.2mm on the tablet (**Figure S2a**). Cursor position was recorded using PsychoPy, with a sampling rate of 60Hz, but was only visible during the task response period.

On each trial, incidental hand motion – approximated by stylus tip position – was recorded from the pre-stimulus fixation until the end of delay. During the recall phase, the tablet and stylus remained active, but recorded only explicit response-related events: drawing, wheel adjustment, and clicking the ‘continue’ button. Participants were instructed to hold the stylus throughout the experiment, but they were otherwise unaware that stylus movements aside from explicit task responses were being recorded.

Note that stylus position was recorded even when it was lifted off the tablet, within a height range of ∼1cm, which is the maximum configurable range for the device. Based on our observations during piloting and practice, the stylus normally remains within this range when participants naturally rest their hand on the table or tablet.

### QUANTIFICATION AND STATISTICAL ANALYSIS

Detailed summary statistics and full statistical outputs are provided in **Data S1**.

#### Recall response processing

Although participants were allowed to redo their response in both response formats, our analyses included only the final response submitted. Because participants made drawings relatively quickly, the movements leading up to and following the intended response might exhibit some artifactual aspects. Indeed, sometimes the beginning and end of the stroke exhibited distortions away from the main axis of recall, so we excluded the portions of each drawing produced during the first and last 10% of the drawing time.

In the *wheel* condition, the response was defined as the orientation of the line connecting the two dots. In the *draw* condition, the response was defined as the average orientation of the strokes remaining after denoising the drawing.

##### Response error

Error was defined as the angular difference between the response and true stimulus orientation. Subject-wise average error was calculated as the mean absolute error. All participants met the behavioral QA criteria (average error <30°).

We quantified memory precision using two complementary metrics. As large errors (> 45°) can disproportionately inflate summary statistics, we first computed subject-wise average error after excluding trials with error >45° (this exclusion was only applied to recall behavior analyses). We also computed the sharpness of each subject’s error distribution around zero, incorporating all trials.

Sharpness was computed by first smoothing the error distribution to address data sparsity:

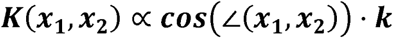

Where ***k*= 20**. Sharpness was then defined as:

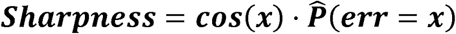

A smaller average error or greater sharpness indicates more precise memory recall.

##### Response time indices

For each trial, we computed two response timing indices. Initiation time was defined as the interval between the onset of the recall phase and the first stylus touch down on the tablet, indexing the speed of response preparation.

Confirmation time was defined as the interval between completing the final response (i.e., the last stylus contact in the response field) and pressing the “continue” button, indexing the speed of post-response verification.

#### Gaze preprocessing and feature extraction

Preprocessing and quality control of gaze data closely followed published procedures^55^. Blinks were identified as time points where the rate of pupil-size change was an outlier (i.e., greater than 4 median absolute deviations). Gaze data from these time points and the immediately neighboring epochs was removed, and the resulting gaps were interpolated. Trials were excluded if more than 10% of data points had been removed, and one participant was excluded from further analysis for having more than 40% of trials excluded. For the remaining participants, 4% of trials were removed for failing the gaze QA.

To characterize gaze patterns, we extracted features as follows. First, we established a time-window-specific baseline for each subject, to address systematic vertical drifts observed across the trial. For each 50 ms window, we calculated the median x and y gaze coordinates across all trials for that time window, for each participant. Gaze data were then re-centered based on this participant- and time-window-specific median location. After re-centering, we analyzed only gaze data falling between 15 and 150 pixels from the normalized center. This was geared to minimize the impact of small vertical displacements (<15 px) that we observed across conditions (maybe driven by a bias toward the tablet position), as well as large jumps (>150 px) that may be attributed to distraction or off-task behavior. But results were unchanged by loosening these thresholds. We then extracted feature vectors for each time window within each trial: the distribution of gaze offset angles, relative to the time-window specific baseline location, within each of 36 directional bins spanning 360° (weighted by magnitude). We also illustrate the spatial distribution of gaze as a two-dimensional heatmap, purely for visualization purposes. To account for subject-specific biases, we computed a baseline feature vector for each subject (the average across all trials, over the task interval from the first stimulus onset to the recall phase onset), and subtracted this baseline from all their feature vectors. To analyze data across-subjects, we applied z-normalization to the resulting feature vectors for each participant, per time window.

#### Hand motion preprocessing and feature extraction

Motions lasting less than 45 ms (i.e., when continuous position changes ceased within 45 ms) were considered sudden jumps and removed as artifacts. The remaining hand motion data were represented as a sequence of motion vectors, each defined by its direction, magnitude, and start and end times. To further smooth the data, two consecutive motion segments with angular differences of less than 15° were merged into a single motion. The direction of the merged motion was computed as the weighted average of the constituent segment motions, and its magnitude was computed as the sum of their individual magnitudes.

Trials with no valid hand movements (defined as movements spanning at least three timepoints at a sampling rate of ∼60HZ) detected between the end of fixation and the start of the response phase were excluded. One participant was excluded from further analysis for retaining fewer than 60% of trials after QA. For the remaining participants, approximately 8% of trials were removed per person; valid hand movements cover approximately 38% of timepoints throughout the delay on average.

To characterize hand motion patterns, we extracted features as follows. For every 50ms time window within a trial, we computed the magnitude of motions within each of 36 directional bins spanning 360° (i.e., 10° per bin), resulting in a 36-dimensional vector. (For motions that spanned multiple time windows, the motion magnitude assigned to each window was proportional to the fraction of the motion’s duration falling within that window). To account for subject-specific biases, we computed a baseline feature vector for each subject (the average across all trials, over the task interval from the first stimulus onset to the recall phase onset), and subtracted this baseline from all their feature vectors. To analyze data across-subjects, we applied z-normalization to the resulting feature vectors for each participant, per time window.

#### Combined QA criteria

Trials were excluded if gaze data failed quality assurance or if no valid hand-motion data were recorded. Unless otherwise specified, all analyses (including recall behavior) include only trials that passed quality assurance for both gaze and hand-motion data (but no gaze or hand data were excluded based on behavioral criteria). As a result, 35 of the 37 subjects who completed the experiment were retained, each with more than 60% of trials remaining after exclusion. Across the retained subjects, ∼12.6% of trials were excluded for failing the quality checks (an average of 19/160 trials per individual).

#### Recall performance statistics

For all response metrics, we used two-sided paired samples t-tests and Wilcoxon signed-rank tests to compare subject-wise metrics between draw and wheel conditions. We analyzed average error (**Figure 1d** left), average initiation time (**Figure 1d** middle), average confirmation time (**Figure 1d** right), and the sharpness of the error distribution (**Figure 1e**).

### Representational similarity analysis (RSA)

#### RSA procedure

We follow a similar approach to that described in previous literature^12^. To quantify similarity between the peripheral motor activity patterns, we use cosine similarity between pattern feature vectors. To quantify similarity between stimuli, we use the absolute angular difference between their orientations.

We calculated gaze or hand motion pattern similarity across trials, stimulus similarity across trials, and the correlation between the two. Specifically, for each participant and each 150 ms sliding window, we considered all pairs of trials ***i*** and ***j*** (***i =t j***), and we constructed two matrices. For each pair, we computed (1) one minus the cosine similarity between the participants’ motor activity feature vectors within that time window (producing the motor activity dissimilarity matrix), and (2) the absolute angular difference between the corresponding stimuli (producing the stimulus dissimilarity matrix). We then calculated the Pearson correlation between the motor activity dissimilarity matrix and the stimulus dissimilarity matrix. Significance was assessed using a two-tailed permutation test (iterations=1000), evaluating whether the observed correlation differed significantly from zero (i.e., no correlation).

#### RSA signatures of remembered vs. dropped stimuli (Figure 2c)

For each stimulus (Stim1 or Stim2), we split trials based on whether the stimulus was cued to be remembered or dropped during presentation. RSA was then applied within each partition.

#### RSA signatures of remembered stimuli under different response format conditions (Figure 3a)

For each stimulus (Stim1 or Stim2), we selected only trials in which the stimulus was cued to be remembered, and then split the trials based on whether it was in the *draw* or *wheel* condition. RSA was then applied within each partition.

### Inverted encoding modeling (IEM)

#### IEM reconstruction procedure

Inverted encoding model (IEM) has often been used to reconstruct an estimate of the perceived or remembered stimuli that are represented across feature-specific information channels, based on aggregate measures of neural activity^31,32^. Here, we adapted this approach to reconstruct the remembered information in gaze and hand motion patterns. IEM takes the features extracted from gaze or hand movements, during a specific task epoch, as input to reconstruct the orientation of the target as a probability distribution across the feature space (i.e., 0-180°). Because the analysis output is distributional, the method is especially well-suited here to characterize the representation of a given WM item when multiple items are maintained simultaneously, and to potentially detect subtle biases in stimulus evidence. Here, we aim to approximate the information that can be detected in aggregate motor activity patterns, but make no claims about the feature-tuning of those signals.

We used a model with 18 channels, each assumed to best respond to a different orientation: 0°, 10°, …, 170°. Let the peripheral signal features be denoted by ***X***, the channel activations by ***C***, and the channels’ basis function as ***W***. Assuming

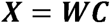

We estimate ***W*** using the peripheral signal patterns ***X***_**1**_ and the estimated channel activations c_1_ from the training data:

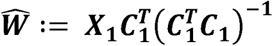

Then, for a separate set of test trials with signal patterns ***X***_**2**_, we can estimate the corresponding channel activations as:

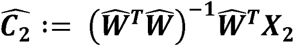

Let ***C***_**z**_ denote the activation of the channel that prefers orientation ***z***. We first define the similarity between two angles (i.e., ***y***_**1**_ and ***y***_**2**_, whose angle between is ***L(y***_**1**_***, y***_**2**_)) as:

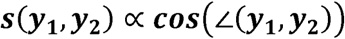

For a given stimulus y, the activation of each channel can be defined as:

We denote the projection from stimulus space ***Y*** to channel activation space ***c*** as π_**Y➔C**_. Thus, based on the predicted channel activations Ĉ, the predicted stimuli ŷ could be obtained by:

The only free parameters in this model are the sharpness parameter k, which determines the tuning width of channel activations. To find the optimal ***k***, we compute the reconstruction loss for all ***k*** as:

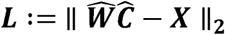

With the optimal sharpness k:

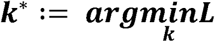

For each trial, the model predicts a distribution, ***̂P(s= x)*** representing the predicted probability that the target stimulus ***s*** has value ***x***. To evaluate model performance, we first compute the distribution of reconstruction errors across all trials for each participant. We then quantify the quality of the reconstruction as follows (similar to previous studies^39^):

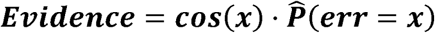

A smaller reconstruction error should correspond to a sharper error distribution around zero, suggesting higher evidence. A uniform distribution of reconstruction errors yields an evidence value of zero. In contrast, a model that systematically tracks the orientation orthogonal to the target is expected to yield a negative evidence value.

Unless otherwise specified, we used a leave-one-subject-out cross-validation scheme: the training set included all trials from all participants except one, and the test set comprised all trials from the held-out participant. When training and testing across different experimental phases or trial conditions, we first split the test data at the subject-level, and then for each test-set selected the training data to ensure no overlap between train and test sets in terms of participants (*across-subjects* model).

Across-subject cross-validation was largely geared to test the generalizability of activity patterns across people. However, it is also an important analysis choice here, to ensure sufficient statistical power given the limited number of trials per subject, per specific condition. While each subject’s trials would only cover a limited set of orientation features in each task condition, a leave-one-subject-out framework yields a training set of hundreds or thousands of trials, fully covering the stimulus space. Combined with other analytic choices such as soft-binning (i.e., applying a Gaussian-kernel to the input-to-channel mappings) and a task design where subjects often remembered multiple items per trial, this should render analyses robust to individual-level data sparsity. In a post-hoc robustness check, both primary gaze- and hand-movement-based IEM analysis results reached a replication rate of 0.8 when trial numbers were downsampled to 80%.

#### IEM evidence across trial epochs (Figure 2d)

IEM analysis could focused on three distinct epochs: (1) post-Stim1 encoding (i.e., the 500ms immediately following the presentation of Stim1); (2) post-Stim2 encoding (i.e., the 500ms immediately following the presentation of Stim2); and (3) the WM delay (i.e., the 5000ms interval following post-Stim2 encoding and before the recall epoch.

Within each epoch, ‘relevant stimuli’ were defined as stimuli had been presented and cued to remember up to that point. For instance, only Stim1 that were cued were considered relevant for the post-Stim1 encoding epoch, whereas both Stim1 and Stim2 that were cued were considered relevant for post-Stim2 encoding and WM delay epochs. For each epoch, using a leave-one-subject-out cross-validation framework, we trained an IEM decoder on data from other participants and tested the reconstruction of relevant stimuli on the held-out subject. The sharpness of the resulting error distribution for Stim1 or Stim2 therefore indexes the strength of ‘evidence’ for that stimulus in that subject (see section above for details about ‘evidence’ metric). We performed one-sided *t*-tests and Wilcoxon signed-rank tests to assess whether the evidence for Stim1 or Stim2 was significantly above chance (i.e., zero) during each epoch. These analyses were conducted independently for gaze and hand-motion-based reconstructions.

#### IEM evidence between response format conditions (Figure 3b)

We used the same delay-epoch reconstructions as described in the previous section (i.e., **Figure 2d**). However, when quantifying reconstruction quality in the test set, we pooled the reconstructions of both Stim1 and Stim2, and grouped them based on whether they were taken from *draw* vs. *wheel* condition trials. Within each subject, the sharpness of the reconstruction error indexes the strength of WM evidence in the subject’s gaze or hand movement patterns, under *draw* or *wheel* conditions. For both gaze-based or hand-based reconstruction, we applied two-sided paired samples *t*-tests and Wilcoxon signed-rank tests to compare the evidence between *draw* and *wheel* conditions.

#### IEM evidence for recalled response (Figure 3c)

We also trained IEMs to reconstruct the orientation of the response that was entered during the recall phase (rather than the encoded stimulus). We followed the same procedures described above for stimulus decoding (using the same leave-one-subject-out framework and delay-period data). We included only stimuli that were cued to be remembered. However, we changed the training and testing targets from the correct presented stimulus values to instead use whatever orientation the subjects reported for recall.

Responses were split into three error subsets based on behavioral error magnitudes. This split was applied after partitioning the data by subject, response format (*draw* vs. *wheel*), and stimulus (Stim1 vs. Stim2), ensuring that responses reflected the relatively best and worst performance within a condition. Within each partition, responses were ranked by error magnitude and the smallest and largest error terciles were designated as the ‘small error’ and ‘large error’ subsets, respectively.

We examined the IEM reconstruction sharpness for each tercile and task condition. We first examined whether behavioral responses were robustly decodable from both gaze and hand movements in both the small and large error terciles. We then applied one-sided paired samples *t*-tests and Wilcoxon signed rank tests to examine whether the evidence for responses was significantly stronger in ‘small error’ (vs. ‘large error’) groupings.

### Gaze biases during selection for recall

During the recall phase, we suppressed collecting stylus data that was not in contact with the tablet. Therefore, for recall phase analyses, we focus on the gaze data (**Figure 2e**). Specifically, we analyze the first 500ms after probe onset, only for trials when both WM items had to be reported. This window is aimed to capture the period when participants would be selecting the recalled item from within WM, but before they execute the motor response (minimum response initiation time is ∼1000ms). We grouped trials based on whether the first recalled item was Stim1 or Stim2. Then, for each subject, we calculated the time series of their average gaze offset (relative to their mean gaze position), as a function of whether Stim1 or Stim2 was recalled first. We applied a cluster-based permutation test to determine if these gaze offsets differed between the two trial groupings.

### Correlated evidence fluctuations across effectors

#### Across subjects (Figure 4c)

To investigate the relationship between modalities across subjects, we performed a linear regression between the gaze-based and hand-based response-format effects (defined as the difference score in IEM evidence between *draw* and *wheel* trials for each subject). We apply a two-sided *t*-test on the regression coefficients. A regression slope significantly below zero suggests a negative relationship between the gaze and hand response-format effects.

#### Within subjects (Figure 4d-e)

We also investigated the coupling between gaze and hand signals within subjects, examining how the evidence fluctuated together both within and across trials (during the delay period). These analyses were geared to further test whether the signals trade-off with each other or co-vary together.

The majority of trials (75%) had a memory load of 2, and we restricted these analyses to those trials. Given that subjects would likely alternate attention between the two items across the trial (**Figure S4a**), we reasoned that the evidence for the currently attended item (rather than the average across both items) may best capture the WM representational quality at any given time point. We therefore first assessed which of the two stimulus signals was ‘dominant’ at each timepoint (i.e., showed greater, non-negative evidence). Specifically, IEM decoders trained on aggregated delay-period data (i.e., **Figure 3b**) were applied to 350ms sliding windows across the delay to reconstruct Stim1 and Stim2 (window size was chosen to balance temporal resolution with signal-noise ratio). Within each window, the item with greater evidence was designated as the ‘dominant’ item, and we took that stimulus evidence as the representational strength at that time point (**Figure S4b**). Therefore, for each trial and effector, we got a time series of representational strength, defined as the time course of maximal evidence (for either Stim1 or Stim2). We then correlated these timecourses between gaze and hand activity.

To assess moment-to-moment interdependence, we quantified the within-trial correlation between gaze- and hand-movement representational strength time series. For each participant, we applied repeated measure correlations (rmcorr^71^; Pingouin implementation^72^), treating each time point as the repeated unit and trial as the grouping factor. Specifically, rmcorr used ANCOVA to estimate a single common regression slope between gaze and hand representational strength across timepoints within each trial’s delay period; the implementation allows each trial to have its own intercept, to account for between-trial differences in mean signal level, yielding a single correlation coefficient per participant. Group-level significance was assessed by testing these subject-level coefficients against zero using two-sided *t*-tests and Wilcoxon signed-rank tests. If the coefficient is significantly above zero, there is a positive correlation between moment-to-moment gaze and hand WM representational strength, while a significantly below zero coefficient suggests the opposite.

To look at across-trial relationships, we then computed the mean representational strength for each trial, across that trial’s delay period time points, yielding one scalar value per trial, per effector. We then applied rmcorr, treating each trial as the repeated unit and participant as the grouping factor; this estimated a common regression slope between trial-averaged gaze and hand representational strength within each individual, while allowing each participant to have their own intercept. We tested whether the estimated slope was significantly different from zero using the p-value derived from the rmcorr model as implemented in Pingouin. A significantly positive (negative) coefficient indicates that gaze and hand WM representational strength are positively (negatively) correlated across trials.

