## Supplemental Information for "Flexible Working Memory in the Peripheral Nervous System"

### SUPPLEMENTAL RESULTS

#### 1. Behavioral precision under different load conditions (Supplementing main text Figure 1)

In the main text, we collapse across trials when one or both working memory (WM) sample stimuli were cued to remember. Here we examine these load conditions separately (**Figure S1a-b**).

We compare behavioral precision metrics between the *draw* and *wheel* response formats as a function of WM load (**Figure S1c-e**). When only one item was remembered, *draw* and *wheel* conditions did not differ on either error ($\boldsymbol{W = 241, n = 35, p = 0.232}$) or sharpness ($\boldsymbol{W = 297, n = 35, p = 0.777}$). When both items were remembered, the *draw* condition showed larger errors ($\boldsymbol{W = 189, n = 35, p = 0.039}$), but no effect on sharpness ($\boldsymbol{W = 267, n = 35, p = 0.441}$).

We also fitted a linear mixed-effects model to predict error magnitude as a function of load, response format, and their interactions. Only the load coefficient differed significantly from zero, while neither the response format nor its interaction with load were significant. Therefore, behavioral error was largely comparable across our different response formats conditions, summarized in the table below (n=35):

**Table S1: Recall error – linear mixed model**

| | Estimate (β) | SE | z | p | 95% CI |
|---|---|---|---|---|---|
| Intercept | 11.01 | 0.53 | 20.97 | <.001 | [9.98, 12.04] |
| Load (2 vs 1) | 1.81 | 0.47 | 3.82 | <.001 | [0.88, 2.73] |
| Format (wheel vs. draw) | -0.56 | 0.47 | -1.18 | 0.238 | [-1.49, 0.37] |
| Load × Format | –0.39 | 0.67 | –0.59 | 0.559 | [-1.70, 0.92] |

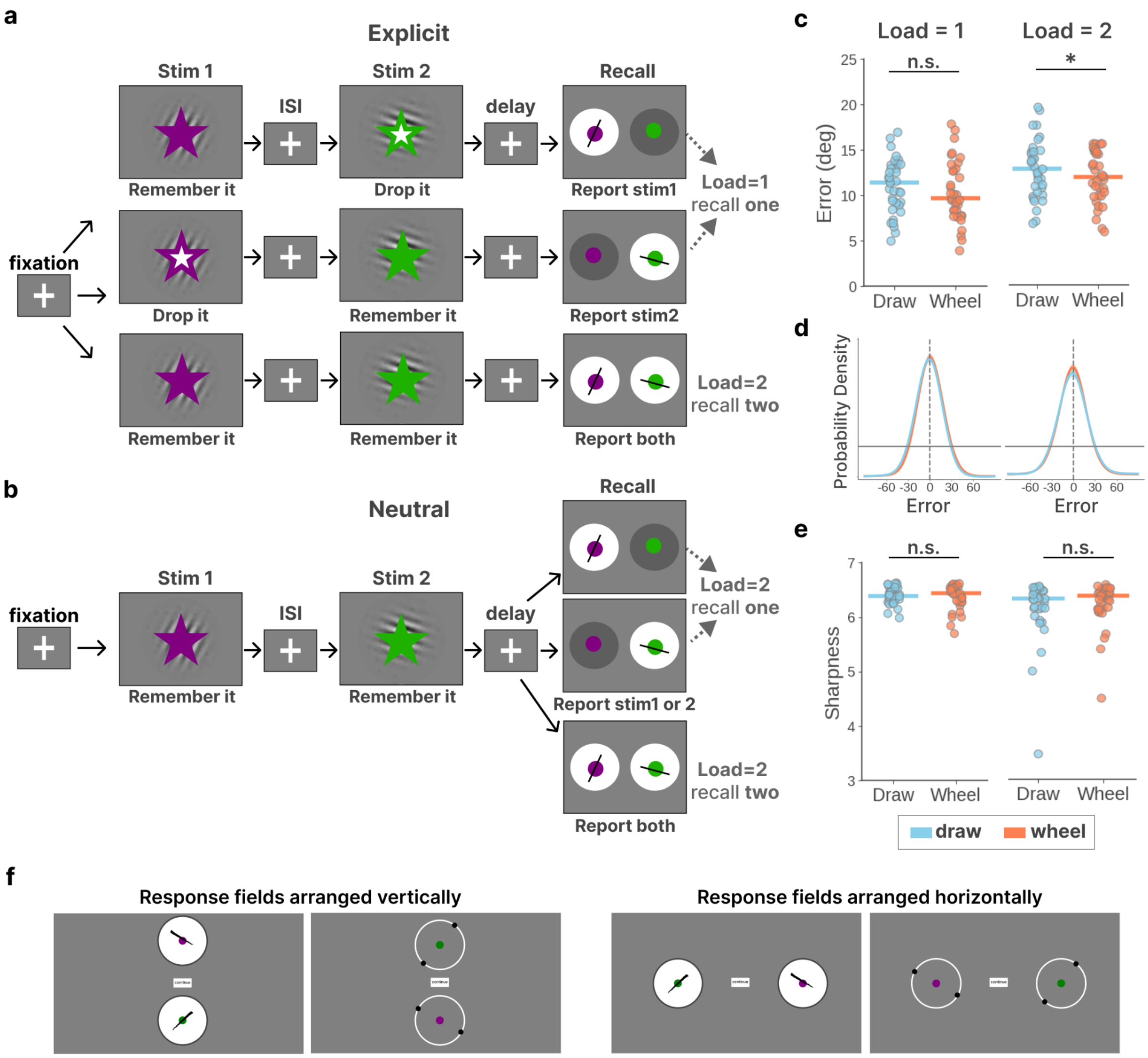


***Figure S1: Supplementing Main text Figure 1.*** *Behavioral performance across load conditions (1 vs. 2).* ***(a-b)*** *Experimental design schematic for the draw condition (the wheel condition used the same design, except the response field entailed a wheel adjustment).* ***(a)*** *In 'explicit' conditions, the cued items were always recalled. When one item was cued, memory load was 1 and one item was reported; when two items were cued, memory load was 2 and both items were reported.* ***(b)*** *In the 'neutral' condition, both items were cued to remember, but participants did not know which would be probed until the response phase. Memory load was always 2; one item was reported on half of trials, and both items were reported on the other half.* ***(c-e)*** *Behavioral performance, for both the draw and wheel conditions, separately for each load level. Left: Load = 1; Right: Load = 2.* ***(c)*** *Mean response*

*error for each individual. **(d)** Distribution of response errors. **(e)** Sharpness of the error distribution curves for each individual. **(f)** Examples illustrating the possible response field arrangements, to scale, where the confirmation button was fixed at screen center.*

---

#### **2. Motor activity characteristics** (Supplementing main text Figure 2)

##### *2.1 Hand movement magnitudes during delay and recall*

Here, we examine the magnitude of hand movements during the delay, to get a sense for their scale relative to explicit drawing responses.

We quantified delay-period hand movements by either their trial-wise span or time-point-wise displacement (**Figure S2b**). Span was defined as the maximum distance between any two stylus locations across the delay on that trial. Displacement was defined as the offset at each timepoint relative to the hand position at the start of the delay. The magnitude of the explicit drawing response was defined as the Euclidean distance between its start and end points. To provide a concrete measure of physical movement, stylus data that were recorded as cursor positions on the screen have been projected onto physical locations on the tablet; thus, an ‘X cm’ movement refers to the stylus moving X cm over the tablet.

Overall, delay period movements were smaller than drawing responses, and the most prevalent delay movements were within the 2mm range, compared to ~4cm for drawings (**Figure S2b**, right). This suggests that delay-period activity preserves the spatial features of the WM content without executing a full motor rehearsal.

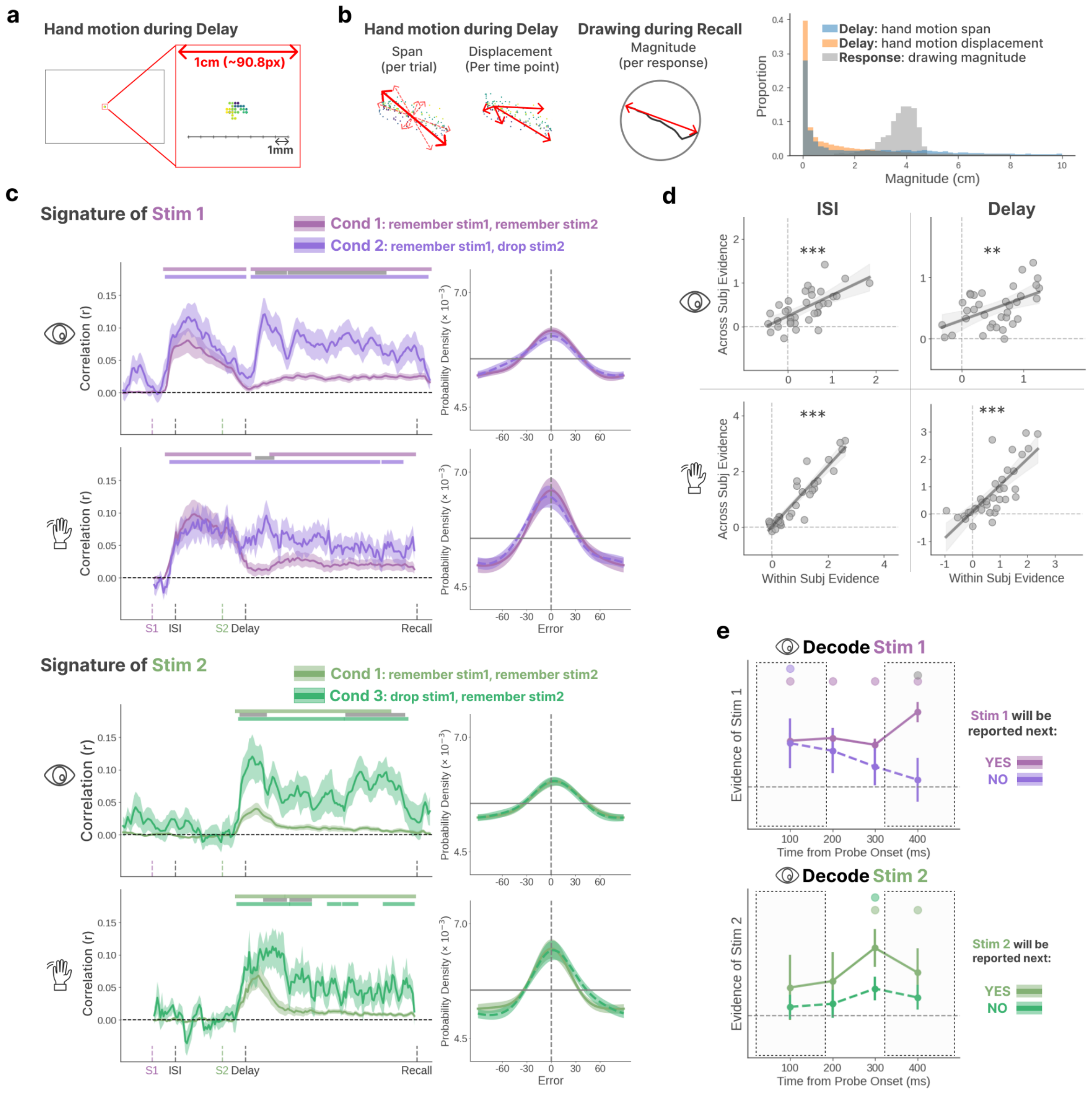


***Figure S2: Supplementing Main text Figure 2. (a)*** *A depiction of the spatial resolution of hand movement recording. A 1cm movement on the tablet corresponds to approximately 90.8 pixel units on the screen. Colored dots are stylus tip positions recorded during the delay period from one trial, with each dot corresponding to the position at a single timepoint.* ***(b)*** *An illustration of how the max span, displacement, and response magnitudes are quantified. Delay period movements are not to scale, relative to the response, as the delay illustrations*

*are relatively enlarged for clarity. The distributions of delay period hand movement and recall period response magnitudes.* ***(c)*** *RSA and IEM reconstructions for gaze and hand motion patterns under different memory load conditions. Evidence for Stim1 and Stim2 are plotted separately for each load condition, to illustrate how the encoding or maintenance of one stimulus depends on whether the other is remembered or dropped.* ***(d)*** *Correlations between within-subjects vs. across-subjects IEM decoding evidence. Shown separately for gaze and hand movement recordings, from both the ISI and delay periods. Within-subjects scores reflect each individual's stimulus reconstruction strength when the model was trained and tested on their own data. Across-subjects scores reflect each individual's stimulus reconstruction strength when the model was trained on the data from all the other subjects and tested on the individual.* ***(e)*** *Stimulus evidence from gaze-based IEM reconstructions during the recall period (the 0.5 s following probe onset). Evidence is plotted separately for each stimulus, depending on whether participants chose to report that stimulus first or second (restricted to trials when both items were cued to remember and report). A sliding window of 200 ms was applied across this period with a step size of 100 ms.*

---

#### 2.2 Stimulus evidence under different load conditions

While two sample stimuli were presented on every trial, sometimes one stimulus was cued to drop. Trials could be classified into three groups depending on which stimuli were cued to be remembered or dropped:

- **Condition 1**: Load 2, both Stim1 and Stim2 were cued to remember. This could occur in 'explicit' blocks when both cued items would be tested, or in 'neutral' blocks when both items were cued but it was uncertain which would be tested (See the **Figure S1a-b** for block design details).
- **Condition 2**: Load 1, only Stim1 was cued to remember while Stim2 was cued to drop ('explicit' blocks).
- **Condition 3**: Load 1, only Stim2 was cued to remember while Stim1 was cued to drop ('explicit' blocks).

The main text results collapsed across these conditions (**Figure 2c-d**). Here we repeat the RSA and IEM analyses, separately for each of the conditions listed above. This will inform how the encoding and maintenance of a given sample item is impacted by whether the other item is remembered or dropped. This will also inform whether these peripheral signatures can code for more than one WM item simultaneously.

As shown in **Figure S2c** left, we detected stimulus evidence for all remembered stimuli, after encoding and throughout the delay, in all conditions. However, for Stim1, its rebound and maintenance after Stim2 encoding were greater when Stim2 could be dropped. Likewise, for Stim 2, its encoding and maintenance were greater when Stim1 could be dropped. However, when we aggregated the signals across the delay period,

IEM reconstruction evidence was comparable across different load conditions (**Figure S2c** right). Notably, the same decoder (i.e., trained on data aggregated across conditions) was used for testing in all conditions, suggesting that the WM representations in these activity patterns were similar regardless of serial encoding position or WM load.

#### 2.3 Generalizing across individuals

In the main text we showed that models trained on data across participants could successfully reconstruct WM stimuli from a held-out participant's data. This implies that the patterns encoding WM features in eye and hand movement activity may generalize across people. In addition to training and testing with a leave-one-subject-out approach (denoted as the 'across-subject' model), we also trained and tested individual models for each subject (denoted as the 'within-subject' model), and examined the relationships between them.

For within-subject analyses, we applied a 5-fold cross-validation scheme, with each fold containing 20% of trials from each participant.

The strength of an individual's stimulus evidence was strongly related between models, as estimated with linear regression slopes (**Figure S2d**; ISI gaze: $\boldsymbol{\beta = 0.48, p < 0.001}$; ISI hand: $\boldsymbol{\beta = 1.09, p < 0.001}$; delay gaze: $\boldsymbol{\beta = 0.36, p = 0.002}$; delay hand: $\boldsymbol{\beta = 0.96, p < 0.001}$). That is, participants whose gaze or hand movement patterns enabled more precise within-subject reconstructions (using models trained on other trials) also showed more precise reconstructions when the model was trained on data from other participants. These findings provide additional evidence for the generalizability of WM-related peripheral activity patterns across individuals.

#### 2.4 Gaze patterns during the WM recall phase

In the main text, we showed horizontal gaze biases during the recall phase, as a function of which sample item was selected to report first (**Figure 2e**). In addition to these directional biases, here we also test the strength of specific stimulus evidence (i.e., orientation) during the recall selection period.

We used IEM reconstruction sharpness to quantify the strength of orientation evidence in gaze patterns (as in **Figure 2d**), but shifted our focus to the 500ms period after probe onset. Instead of using the delay-period-trained decoders, we used a sliding window approach (200ms windows, 100ms step) throughout the post-probe period, training IEM decoders for each time window separately. Because the gaze biases for Stim1 and Stim2 drifted in opposite directions, we trained separate decoders for each stimulus within each time window. Therefore, for each subject, we trained a separate decoder for each time window and each stimulus, using data pooled from all other subjects.

We limited this analysis to trials where participants were cued and probed to report both items. On these trials, participants were free to choose which of the two items they wanted to report first. Thus, the first 500ms after probe onset, which preceded the explicit response, should index the volitional selection process within WM. We grouped trials based on whether Stim1 or Stim2 was reported first, and we compared the stimulus evidence in gaze patterns, across the selection period, for items that were reported first or second.

As shown in **Figure S2e**, stimulus evidence diverged based on reporting order. When an item was selected to report first, its code was sustained (Stim2) or strengthened (Stim1) throughout the selection period. In contrast, when the same item was not selected for immediate report (despite remaining relevant for the future), its representation was relatively weaker overall (Stim2) or progressively dampened (Stim1). We should note that participants exhibited a strong preference for reporting Stim1 first ($\mathrm{M}{=}76\%$ of trials), which was significantly above the 50% chance level ($p{<}.001$), consistent with a primacy effect. However, the evidence difference between the immediate and deferred conditions became more pronounced over the course of the selection period for both Stim1 and Stim2. This result shows how gaze activity is engaged not only during maintenance, but also during stimulus selection for a manual response.

#### 3. Characteristics of condition-specific effector preferences (Supplementing main text Figure 3)

##### *3.1 Generalizing across response formats*

In the main text, we reported successful reconstruction of the memorized orientation from gaze and hand motion patterns using a decoder trained on data pooled from both *draw* and *wheel* conditions. This was based on our assumption that the encoding of WM features in peripheral motor signals (i.e., the mapping between memorized orientation and gaze/hand movement patterns) would be consistent across response formats, even if the strength of that encoding could differ (as shown in main text). In other words, we assume that if people tend to look more rightward when thinking of 45° in the *draw* condition, a similar rightward bias should also be present for 45° in the wheel condition – even if the precise mean position, variance, or strength of the bias differs. Thus, we assumed that our response format manipulation should not qualitatively change how WM features are encoded in peripheral signals, but may induce a quantitative change in the strength of stimulus evidence.

Here, we provide supplementary analyses to support this assumption. First, for trials with a memory load of 1, we grouped trials by their memorized orientation (as in **Figure 2a**) and plotted the normalized patterns of gaze and hand motions during the delay separately for each response format (**Figure S3b**). Visually, the gaze and hand motion patterns appear descriptively similar across the two conditions. For instance, we observed greater hand movements in the direction of the memorized orientation, and a rotation of the gaze 'hotspot' as the memorized orientation rotated.

To quantify this apparent similarity, we trained and tested the IEM model on data from each response format condition separately, and compared the reconstruction errors both within- and across-formats (**Figure S3c**). The reconstruction results of the model trained on format A and tested on format B is denoted as 'A → B'. If the encoding patterns differ qualitatively between response format conditions, we would expect cross-format decoding (i.e. 'A → B') to be significantly worse than within-format (i.e. 'A → A' and 'B → B'). The results are summarized in the table below.

**Table S2: IEM evidence cross- vs. within-formats**

| | cross-formats | within-formats | $t(34)$ $(p)$ | $p$(normal) | $W(n = 35)$ $(p)$ |
|---|---|---|---|---|---|
| Gaze | draw → wheel | draw → draw | 1.22 (0.232) | 0.727 | 235 (0.195) |
| | | wheel → wheel | 1.16 (0.254) | 0.968 | 252 (0.310) |
| | wheel → draw | draw → draw | -0.99 (0.330) | 0.484 | 259 (0.368) |
| | | wheel → wheel | -1.37 (0.180) | 0.311 | 243 (0.245) |
| Hand | draw → wheel | draw → draw | -4.05 (<.001) | 0.289 | 107 (<.001) |
| | | wheel → wheel | -0.11 (0.917) | 0.565 | 293 (0.728) |
| | wheel → draw | draw → draw | 1.21 (0.234) | 0.452 | 261 (0.385) |
| | | wheel → wheel | 4.99 (<.001) | 0.041 | 75 (<.001) |

With the significance threshold adjusted using Bonferroni correction ($\mathbf{0.0125}$), we found no significant difference between cross-format and within-format comparisons in the gaze data. In the hand data, ‘*draw* → *draw*’ shows significantly better reconstruction than ‘*draw* → *wheel*’, while ‘*wheel* → *wheel*’ shows significantly worse reconstruction than ‘*wheel* → *draw*’. Rather than showing any advantage of within-format decoding over across-format decoding, these results indicate only a response-format superiority: hand-based decoding evidence is stronger when the test condition is ‘*draw*’ than ‘*wheel*’, regardless of the condition on which the decoder was trained.

As we did not find evidence that within-format decoding is superior, we assume that response format does not qualitatively alter peripheral WM coding.

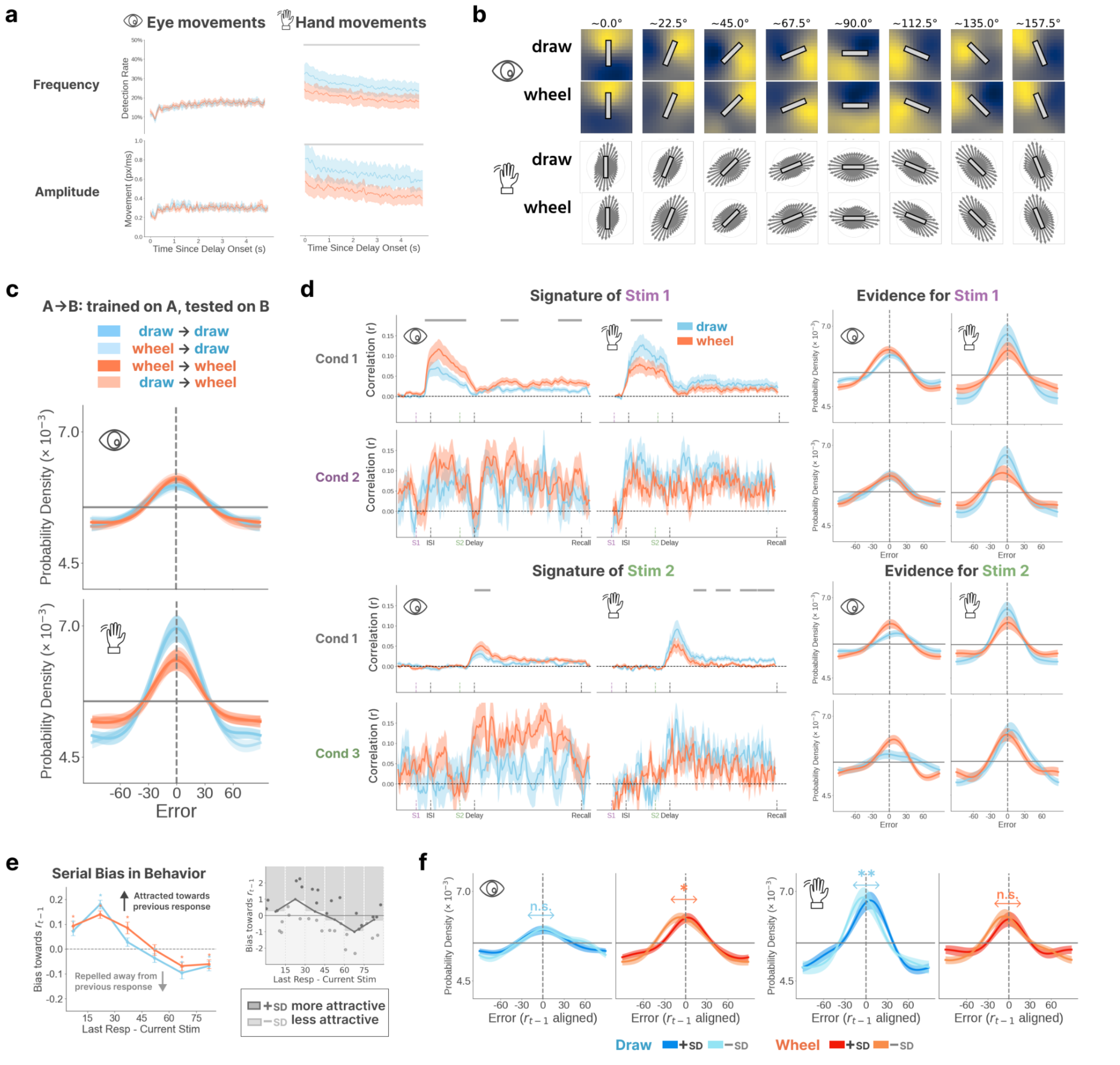


***Figure S3: The impact of response format emerges faster in gaze than in hand motions.*** ***(a)*** *Frequency and magnitude of eye and hand movements throughout the delay period. Event detection threshold is set to 8°/s (or 269px/s).* ***(b)*** *Gaze patterns (i.e. normalized 2D heatmaps of gaze distribution) and hand motion patterns (i.e. normalized angular distribution) as shown in Figure 2a, are plotted separately for the draw and wheel condition.* ***(c)*** *Distributions of reconstruction errors from the IEM analysis using a leave-one-subject-out cross validation paradigm. Each line corresponds to training the model on gaze or hand motion data from either the draw or wheel condition and tested on one of the two conditions.*

*Error distribution curve for the model trained on condition A and tested on condition B is denoted as 'A → B'.* ***(d)*** *Representational similarity timecourse, IEM reconstruction and subjects' IEM evidence for WM items under different load and response format conditions (blue for draw and red for wheel condition).* ***Top:*** *For the first item (Stim1).* ***Bottom:*** *For the second item (Stim 2).* ***(e)*** *Serial dependence in behavior and the diagram illustrating how trials are further grouped based on behavior to examine aggregate stimulus reconstructions. The serial bias could be visualized by plotting the folded signed error as a function of the angular difference between the previous response and the current stimulus (grouped into six bins). For each subject, trials are first binned by the angular difference between the current target and the previous response (six bins). Within each bin, trials in the upper half of values for that bin are considered relatively more attractive ('+ group'), while trials in the lower half for that bin are considered relatively less attractive ('− group').* ***(f)*** *IEM reconstructions (error distributions of model predictions) from gaze and hand motion data, grouped according to whether the behavioral responses on that trial were relatively more or less attractively biased (as described in panel d). Reconstructions are shown for the preferred task condition for each motor effector (i.e., wheel condition for gaze, draw condition for hand motions). The distributions are combined across trials and aligned such that positive values (rightward shift) reflect an attractive bias toward the previous trial, and negative values (leftward shift) reflect a repulsive bias away from the previous trial.*

#### 3.2 Impact of WM load on condition-specific effects

In the main text, we analyze the response-format effects (*draw* vs. *wheel*) by aggregating signals across all cued stimuli (**Figure 3a-b**). Here, we repeat the analysis for each stimulus separately, within each of the three conditions described in ***Stimulus evidence under different load conditions***. Note, however, that Cond 1 (where load = 2) was much more frequent than either Cond 2 or 3 (where load = 1). Therefore, Conds 2 and 3 are likely insufficiently powered here, and the *draw* vs. *wheel* difference observed in the main text is no longer significant, although the descriptive trend remains (**Figure S3d**). However, the main results replicate for Cond 1, when both stimuli are cued to remember, showing a *draw* benefit in hand-based evidence and a *wheel* benefit in gaze-based evidence. These results further suggest that peripheral motor activity can code for more than one WM item simultaneously, with representational evidence being shifted by expected action demand.

**Table S3: IEM evidence – Draw vs. Wheel**

| | M $\pm$ SEM | $t(34)$ $(p)$ | $p$(normal) | Mdn (Q1, Q3) | $W(n = 35)$ $(p)$ |
|---|---|---|---|---|---|
| **Cond 1: Remember both stimuli** | | | | | |
| Gaze | -0.26 $\pm$ 0.09 | -2.75 (0.009) | 0.523 | -0.27 (-0.62, 0.10) | 167 (0.014) |
| Hand | +0.42 $\pm$ 0.12 | +3.61 (0.001) | 0.143 | +0.22 (-0.08, 0.91) | 125 (0.001) |
| **Cond 2: Remember Stim1** | | | | | |
| Gaze | -0.02 $\pm$ 0.19 | -0.12 (0.905) | 0.556 | -0.11 (-0.58, 0.42) | 313 (0.981) |
| Hand | +0.36 $\pm$ 0.19 | +1.95 (0.059) | 0.757 | +0.37 (-0.21, 1.15) | 187 (0.036) |
| **Cond 3: Remember Stim2** | | | | | |
| Gaze | -0.33 $\pm$ 0.14 | -2.27 (0.029) | 0.733 | -0.38 (-0.87, 0.34) | 181 (0.027) |
| Hand | +0.17 $\pm$ 0.18 | +0.93 (0.358) | 0.434 | +0.27 (-0.38, 0.77) | 247 (0.273) |

#### 3.3 Serial dependence in motor activity patterns

In addition to the motor response context (i.e., **Fig. 3**), events of the recent past can also influence WM performance and cortical representations[1–4]. If the observed peripheral WM signatures are more than fleeting motor traces, but instead adaptively anticipate task-relevant behavior, they should also incorporate such temporal context. Therefore, we also tested whether gaze and hand movement WM signatures exhibit serial dependence, and how it corresponds to behavior.

Serial dependence describes a phenomenon whereby WM reports on a given task trial tend to be biased in the feature direction of a previous stimulus or response, as long as the two are fairly similar[5–7]. For instance, a red object may be seen as more orange if you just saw a similar yellow object. This bias is thought to reflect an adaptive temporal smoothing, to strengthen perception and memory in noisy environments[8–10]. If that is the case, we would also expect serial dependence effects to be maximal in the more task-relevant peripheral motor signatures.

Consistent with prior findings[11–13], we observed an attractive bias in behavioral error when the previous response was similar to the current stimulus (**Figure S3e** left). We then tested whether this behavioral bias corresponded with biased stimulus evidence in the peripheral motor activity patterns.

To quantify whether preceding trials impacted current trial peripheral motor signals, we measured whether the delay-period IEM reconstructions were shifted off-center either toward or away from the previous trial's response[14]. For each trial, we aligned the IEM error distributions according to the relationship between current and previous trials: if the last response on the previous trial was counter clockwise relative to the current target stimulus, the error distribution was flipped at zero. The serial bias of the aggregated distribution is defined as follows:

$$\boldsymbol{Bias} = \boldsymbol{sin}(\boldsymbol{x}) \cdot \widehat{\boldsymbol{P}}(\boldsymbol{err} = \boldsymbol{x})$$

We grouped trials by whether behavioral reports were relatively more or less biased toward the previous response ('+SD' group vs. '−SD' group; **Figure S3e** right). We sorted the IEM error distributions, for gaze- and hand-based activity, according to these groups and the *draw* vs. *wheel* task conditions. A positive bias indicates that the evidence is shifted toward the previous response, suggesting an attractive serial effect, while a negative bias indicates a shift away from the previous response, suggesting a repulsive serial effect. We applied paired *t*-test and Wilcoxon signed-rank tests to assess whether the bias was more attractive in the '+SD' group than in '−SD' group.

For both gaze and hand data, trials that showed a more attractive behavioral bias also showed stimulus reconstructions that were more strongly shifted toward previous trial reports (**Figure S3f**), but only for the task conditions that are 'preferred' by each peripheral motor effector (i.e., *wheel for* gaze, *draw for* hand):

**Table S4: Behavior-relevant serial bias in peripheral signals**

| | Condition | M ± SEM (×0.01) | $t(34)$ $(p)$ | $p$(normal) | Mdn (Q1, Q3) (×0.01) | $W(n = 35)$ $(p)$ |
|---|---|---|---|---|---|---|
| Gaze | draw | 0.05 ± 0.19 | 0.25 (0.402) | 0.129 | -0.31 (-0.84, 0.93) | 315 (0.503) |
| | wheel | 0.40 ± 0.23 | 1.78 (0.042) | 0.131 | 0.40 (-0.63, 1.07) | 419 (0.045) |
| Hand | draw | 0.84 ± 0.26 | 3.24 (0.001) | 0.088 | 0.66 (-0.20, 1.96) | 483 (0.003) |
| | wheel | 0.36 ± 0.24 | 1.49 (0.072) | 0.474 | 0.33 (-0.45, 1.28) | 397 (0.092) |

In summary, during WM maintenance, stimulus information in eye and hand-movement signatures exhibited behavior-related serial dependence in 'preferred' task conditions. Signals in the peripheral motor system that are sensitive to prospective task demands may also reflect the integration of perceptual and decisional information over time.

#### 4. State-switching dynamics during maintenance of two items (Supplementing main text Figure 4)

While RSA allows us to examine the aggregate time course of representational information, the approach necessitates combining across trials to estimate the representational geometry of the feature space. In contrast, a sliding-window IEM approach can examine a representational timecourse at the single-trial level, allowing us to examine potential trade-offs between WM items and motor effectors at a fine temporal scale.

Complementing **Figure 4d-e**, here we illustrate the analyses that went into determining which of the two WM sample items was in the 'dominant' state at any given time point. For each peripheral signal within a given time window, the posterior of Stim1 or Stim2 being relatively prioritized was estimated based on its IEM evidence relative to the other item (**Figure S4a**). To visualize the tendency for subjects to alternate between stimuli, we aggregated these posterior time series across trials and clustered them via hierarchical agglomerative clustering (using 1 minus correlation as the distance metric). To ensure the two WM items (and associated states) could be reliably differentiated, these analyses included only trials where the two stimuli were at least 30° apart. Transitions between items appear to occur on the order of seconds (**Figure S4a**, rightmost raster plots after clustering). These state timeseries were then correlated as described in ***Correlated evidence fluctuations across effectors.***

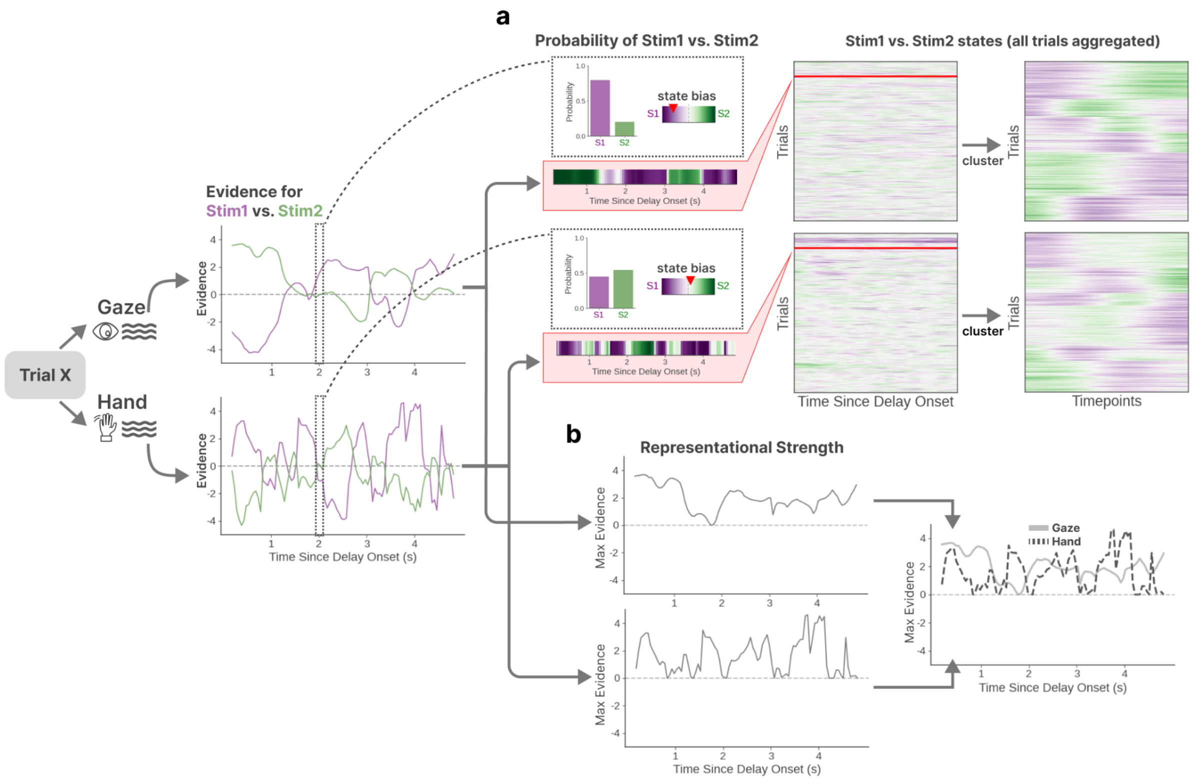


***Figure S4: State of oscillation estimated from IEM decoding.*** *An example trial illustrating the derivation of representational strength for gaze and hand-motion signals during the delay period. Time courses of IEM evidence are calculated independently for Stim1 (purple) and Stimulus 2 (green) for both modalities.* ***(a)*** *State estimation and trial clustering. For a given trial, at each timepoint, the posterior for Stim1 or Stim2 is computed by applying a softmax function to the rectified (non-zero) evidence. State-decoding results are color-coded to indicate a bias toward Stim1 (purple) or Stim2 (green). For each modality, the decoded posterior time series are aggregated across all trials and clustered based on their temporal correlations.* ***(b)*** *Time courses of representational strength. At each timepoint, representational strength is defined as the maximum evidence between Stim1 and Stim2. The resulting time series for gaze (light solid) and hand-motion (dark dashed) are then aligned and correlated.*
